# Evolution of multiple postzygotic barriers between species of the *Mimulus tilingii* complex

**DOI:** 10.1101/2020.08.07.241489

**Authors:** Gabrielle D. Sandstedt, Carrie A. Wu, Andrea L. Sweigart

**Affiliations:** Department of Genetics, University of Georgia, Athens GA, 30602, USA; Department of Biology, University of Richmond, Richmond VA, 23173, USA

**Author notes:** **Corresponding author**: Gabrielle D. Sandstedt, Department of Genetics, University of Georgia, 120 E. Green St. Athens, GA 30602.

**Keywords:** Speciation, Allopatric, Reproductive Isolation, *Mimulus*

## Abstract

Species are often defined by their ability to interbreed (i.e., Biological Species Concept), but determining how and why reproductive isolation arises between new species can be challenging. In the *Mimulus tilingii* species complex, three species (*M. caespitosa*, *M. minor*, and *M. tilingii*) are largely allopatric and grow exclusively at high elevations (>2000m). The extent to which geographic separation has shaped patterns of divergence among the species is not well understood. In this study, we determined that the three species are morphologically and genetically distinct, yet recently diverged (<400kya). Additionally, we performed reciprocal crosses within and between the species and identified several strong postzygotic reproductive barriers, including hybrid seed inviability, F1 hybrid necrosis, and F1 hybrid male and female sterility. In this study, such postzygotic barriers are so strong that a cross between any species pair in the *M. tilingii* complex would cause nearly complete reproductive isolation. We consider how geographical and topographical patterns may have facilitated the evolution of several postzygotic barriers and contributed to speciation of closely related members within the *M. tilingii* species complex.

## INTRODUCTION

Since Darwin initially proposed that natural selection commonly drives the origin of species (Darwin 1859), evolutionary biologists have investigated fundamental questions of how and why new species evolve. Theory suggests that most speciation events begin in allopatry, where geographical barriers prevent gene flow and allow populations to diverge ecologically and genetically (Coyne and Orr 2004). As a byproduct of this divergence, reproductive isolation arises between incipient, sexually reproducing species due to prezygotic barriers that prevent fertilization (e.g., differences in mating system, reproductive timing, or behavior) or postzygotic barriers that cause low fitness in hybrids (e.g., inviability and sterility). Following secondary contact, multiple barriers often act in concert to limit genetic exchange between species (Schluter 2001, Rieseberg and Willis 2007), and selection against hybrids can give rise to additional barriers that further enhance reproductive isolation (i.e., reinforcement; Dobzhansky 1951, Butlin 1989). Because species are often defined by their potential to interbreed (Biological Species Concept, Mayr 1942), a major goal of speciation research is to determine which reproductive barriers evolve during the initial stages of divergence.

One common approach to this problem has been to quantify the relative contributions of pre- and postzygotic barriers to total reproductive isolation (e.g., Ramsey et al. 2003). In plants, such studies often find that prezygotic isolation is stronger than postzygotic isolation (Lowry et al. 2008, Baack et al. 2015), and because reproductive isolating mechanisms act sequentially, it has been argued that the role of later-acting postzygotic barriers is diminished even further (Ramsey et al. 2003, Sobel et al. 2010). Nevertheless, it is important to note that current estimates of reproductive isolating barriers might not reflect their historical roles in species divergence (Widmer et al. 2009), and early-acting barriers can mask later-acting ones, regardless of the order in which they evolved. It is also clear that plant lineages show tremendous variation in patterns of reproductive isolation (Baack et al. 2015) and, in some cases, postzygotic barriers can be quite strong (e.g., Lowry et al. 2008, Ishizaki et al. 2013, Suni and Hopkins 2018, Ostevik et al. 2016, Christie and Strauss 2019).

The extent of geographic overlap between diverging species might also influence the relative importance of pre-versus postzygotic isolation in a given species pair. The potential for increased prezygotic isolation in sympatry due to reinforcement is well established (Coyne and Orr 1989, Noor 1999, Hopkins 2013), but the effect of geography on postzygotic isolation has received less attention. Although intrinsic postzygotic isolation between plant species can be due to chromosomal rearrangements (specifically, hybrid sterility: Stebbins 1958, Rieseberg 2001), in most cases, it is caused by genic incompatibilities (Fishman and Sweigart 2018). When species occur in complete allopatry, these incompatible alleles can evolve to high frequency – either by natural selection or genetic drift – because their negative effects are never exposed in hybrid genomes (Dobzhansky 1937, Muller 1942). In contrast, for species with ongoing gene flow, the maintenance of incompatibility alleles *requires* that their benefits within species outweigh the costs of producing sterile or inviable hybrids (Bank et al. 2012). Thus, neutral or weakly selected incompatibility alleles that might evolve readily in allopatry are expected to be purged from species with extensive geographic overlap and hybridization. Given that many closely related plant species are connected by at least moderate levels of gene flow (Morjan and Rieseberg 2004), this geographic discrepancy might help explain the somewhat lower prevalence of intrinsic postzygotic isolation in plants.

In this study, we investigate reproductive isolation in the *Mimulus tilingii* species complex, a group of yellow mountain monkeyflowers restricted to high elevations (>2000m) along alpine and subalpine streams in western North America (Grant 1924, Pennell 1951). Recently, the *M. tilingii* complex was subdivided into three morphological species – *M. tilingii*, *M. caespitosa*, and *M. minor* – that appear to be largely allopatric (Nesom 2012, 2014). First, we evaluate whether these putative species remain morphologically distinct when grown in a common environment and whether they show evidence of genetic differentiation. Next, we quantify several potential post-pollination barriers by performing reciprocal crosses within and between the three putative species. Surprisingly, we find multiple strong postzygotic isolating barriers between these *M. tilingii* complex species. We argue that strict allopatry might have facilitated the evolution of hybrid incompatibilities in this system, resulting in exceptionally strong postzygotic isolation.

## MATERIALS AND METHODS

### Study System

Members of the *Mimulus tilingii* complex are mat-forming perennials restricted to high elevations west of the Rocky Mountains (Grant 1924, Pennell 1951). They are self-compatible but thought to be predominantly outcrossing, with large, bee-pollinated flowers. We note that some taxonomists have recently reclassified several *Mimulus* species as *Erythranthe* (including the Tilingii group in Nesom 2012, 2014), but because there is still much debate on the status of these taxa (Lowry et al. 2019, Nesom et al. 2019), we continue to refer to them here as *Mimulus*.

The taxonomic status of species in the *M. tilingii* complex has changed over the years, as taxonomists have attempted to grapple with the rich morphological and ecological diversity of the yellow monkeyflowers. In 1974, Vickery informally identified four distinct entities of *M. tilingii*: *M. tilingii* var. *tilingii* (Regel), *M. tilingii* var. *corallinus* (Greene)*, M. implexus* (Greene), and *M. caespitosus*. Through crossing studies, he discovered multiple strong reproductive isolating barriers between *M. t.* var. *tilingii* and *M. t.* var. *corallinus* (including low seed production, low seed germination, and strong F1 sterility; Vickery 1974), but these barriers were largely attributed to differences in chromosome number (n = 14 in *M. t.* var. *tilingii* and n = 24-28 in *M. t.* var. *corallinus*: Mukherjee and Vickery 1959, 1960, and 1962). In the last decade, Nesom (2012, 2013, 2014, 2019) has suggested several species revisions of the *M. tilingii* complex on the basis of differences in morphology (measured in the field and herbaria) and geographic location. Although his species designations vary somewhat among treatments, most include *M. tilingii*, *M. caespitosa*, and *M. minor* with the first two analogous to Vickery’s designations. Of the three putative species, *M. tilingii* is the most widespread, growing throughout much of western North America, whereas *M. caespitosa* grows only in Washington state and southwest Canada, and *M. minor* is restricted to Colorado (Nesom 2012).

### Plant material and care

When we began this study, we tentatively classified plants from 12 populations (17 maternal families) within the *M. tilingii* species complex into three putative species: *M. caespitosa*, *M. minor*, and *M. tilingii*. These 12 populations are distributed across the geographic range of the *M. tilingii* complex (Figure 1; Table S1) and putative species assignments were based on population location (Nesom 2012).

**Figure 1.**
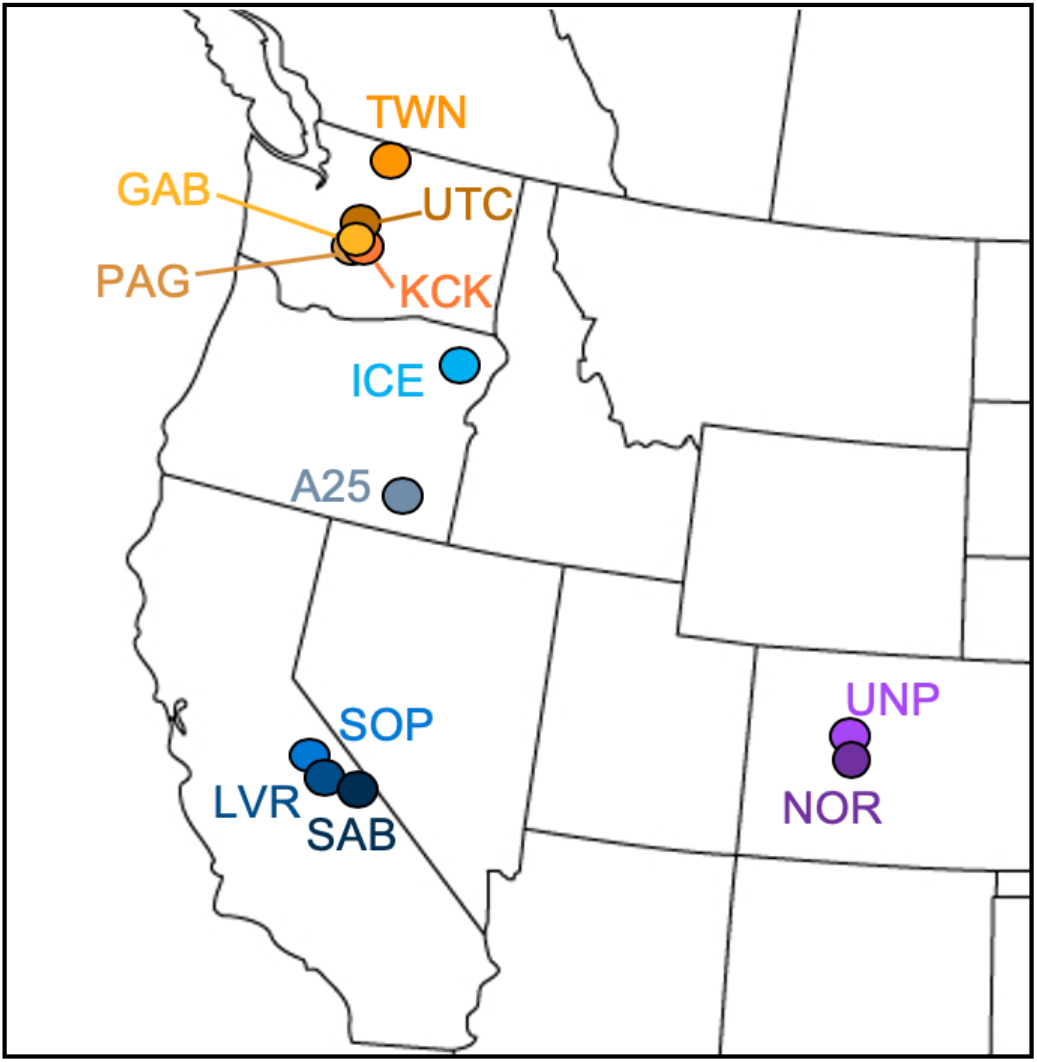
Distribution map of samples in the *Mimulus tilingii* species complex used in this study. Population identity is indicated by a three-letter population code and color. *Mimulus caespitosa* populations are colored in shades of orange, *M. minor* populations are colored in shades of purple, and *M. tilingii* populations are colored in shades of blue.

All maternal families were self-fertilized for one to eight generations (excluding A25; Table S1). To generate experimental plants, seeds were sown onto wet paper towels in petri dishes, sealed with parafilm, and cold-stratified at 4°C for seven days to disrupt seed dormancy. After cold-stratification, petri dishes were transferred to a growth chamber that provided constant supplemental light at 26°C. After germination, seedlings were transplanted to 3.5” pots with moist Fafard 4P growing mix (Sun Gro Horticulture, Agawam, Massachusetts, USA) and transferred to a growth chamber with 16h days at 23°C and 8h nights at 16°C. For assessments of hybrid plant viability and fertility, seedlings were allowed to establish in the growth chamber then moved to a 16h, 23°C/8h, 16°C greenhouse.

### Measuring morphological variation

To characterize genetically-based morphological differences among species within the *M. tilingii* complex, we grew 66 plants from 11 populations (16 maternal families) together in a growth chamber (Table S1). We measured a suite of 16 floral and vegetative traits (Figure S1, Table S2). First, we measured four leaf traits: when the third leaf pair was fully expanded, we used one leaf from the second leaf pair to measure leaf length and width, petiole length, and number of trichomes that exerted past the edge of the leaf (then standardized by leaf length). Next, we measured ten flower traits from one flower on the second flowering pair: corolla height and width, corolla tube length and width, stamen length, pistil length, pedicel length, capsule length, calyx length, and degree of flower nodding. When performing floral measurements, we also measured two stolon traits: number of stolons and stolon length. All traits were measured using calipers, except for the degree of flower nodding, which was measured on photographs using imageJ (Rasband 1997).

We assessed morphological differentiation among species in the *M. tilingii* complex by performing a linear discriminant analysis (LDA), which maximizes variance between predetermined classes (in our analyses, these corresponded to the three putative species) and projects those differences onto a two-dimensional subset. When needed, we transformed morphological trait values to meet LDA assumptions (i.e., that values are normally distributed and means are centered and scaled to zero, Table S2). We assigned each of the 66 plants to *M. caespitosa*, *M. minor*, or *M. tilingii.* To model the LDA, we used the lda function in the R package ‘MASS’. For samples with missing values (traits that were missing or not measured), we used the R package ‘mice’ to impute missing data using a predictive mean matching (PMM) method with 50 iterations. We produced 95% confidence intervals with the R package ‘ellipse’. Finally, we used the R package ‘caret’ and predict function to determine the probability that proposed species in the *M. tilingii* complex correspond to classes predicted by the LDA model.

### Determining genetic diversity and divergence

We generated whole genome sequence (WGS) data for 14 individuals used in the morphology assessments of the previous section (seven *M. caespitosa,* three *M. minor,* four *M. tilingii*). These 14 individuals were from distinct maternal families collected from nine populations (Table S1). We extracted DNA from bud and leaf tissue using a standard CTAB-chloroform protocol (Doyle and Doyle 1987). We submitted the 14 DNA samples to the Duke Center for Genomic and Computational Biology (GCB), which prepared standard 500-bp DNA-seq libraries and sequenced them on the Illumina HiSeq 4000 platform to produce 150-bp paired-end reads. In addition to these newly generated WGS data, we used existing data from two *M. tilingii* samples (A25 and LVR; Garner et al. 2016, Table S1). For population genomic comparisons, we also used existing WGS data for four *M. guttatus*, two *M. nasutus*, and one *M. dentilobus* individuals (Table S3; Brandvain et al. 2014).

To process sequence data, we first trimmed adapters and low-quality bases using Trimmomatic (Bolger et al. 2014) and confirmed removal using FastQC (Andrews 2010). Next, we aligned trimmed paired-end reads to the *M. guttatus* v2.0 unmasked reference genome (http://www.phytozome.net) using BWA-MEM (Li 2013, Li and Durbin 2009). To filter the initial alignment, we used the view command in SAMtools to remove reads with an alignment quality below Q29 (Li et al. 2009). We processed the alignments using Picard tools (http://broadinstitute.github.io/picard); we added read groups with AddorReplaceReadGroups and removed potential PCR and optical duplicates with MarkDuplicates. To confirm paired-end reads mapped together, we used SAMtools fixmate and view commands. To produce a set of high-quality invariant and variant sites for all lines, we used Genome Analysis Toolkit’s (GATK) HaplotypeCaller and performed joint genotyping using GenotypeGVCFs (McKenna et al. 2010). Subsequent filtering and analyses were performed using reference scaffolds 1-14 that correspond to the 14 chromosomes in the *Mimulus* genome. To obtain high-quality genotypes, we used GATK’s VariantFiltration tool to apply hard filtering to sites with mapping quality (MQ) below 40, mapping quality rank sum (MQRankSum) below −12.5, fisher strand (FS) above 60, quality depth (QD) below 2, and read position rank sum (ReadPosRankSum) below −8. We further filtered all sites by removing indels using GATK’s SelectVariants. For each sample, we set a minimum depth of at least ten reads per site and a maximum depth of two standard deviations above the mean read depth, which was calculated after the initial alignment using Qualimap2 (Okonechnikov et al. 2015). We restricted all polymorphic sites to biallelic and, for heterozygous sites, randomly assigned one of the two alleles. Note that because most samples were naturally or artificially inbred, individual heterozygosity was generally low (0.44% - 2.06%; Table S1). Finally, we extracted fourfold degenerate sites from each sample (using a script courtesy of Tim Sackton: https://github.com/tsackton/linked-selection/tree/master/misc_scripts).

To examine patterns of genomic variation in the *M. tilingii* complex, we used a VCF that contained polymorphic (SNP), fourfold degenerate synonymous sites. We selected sites with more than one copy of the minor allele and genotypes for at least 17 of the 21 samples. Note that sample A25 was excluded from these analyses because it had much lower sequence coverage than all other samples with < 20% coverage at these sites. We down-sampled this polymorphic VCF by randomly selecting 1000 SNPs per chromosome, totaling 14,000 sites. We characterized genetic differentiation among individuals in the *M. tilingii* complex using a neighbor-joining (nj) tree. To produce a nj tree, we first converted our SNP genotype file to a pairwise distance matrix. Then, we used the nj function in the R package ‘ape’ to construct a nj tree rooted by the outgroup *M. dentilobus* and rate-smoothed using the function chronopl, where *λ* = 1 (Paradis et al. 2004). We produced a list of 1000 bootstrapped trees using the package ‘phangorn’ and plotted the distribution of trees using Densitree (Schliep 2010, Bouckaert 2010). We also explored genetic relatedness among species in the *M. tilingii* complex using a principal component analysis (PCA). To perform this PCA, we used the function pca in the R package ‘SNPRelate’ and plotted the first two principal components using the R command plot to visualize genetic clusters (Zheng 2013).

In addition to these analyses to visualize genomic structure, we used a VCF containing monomorphic and polymorphic genotype calls at fourfold degenerate sites to calculate pairwise sequence diversity (*π*_*s*_) and divergence (*d*_*s*_). To perform these calculations, we used a python script (Notes S1 in Garner et al. 2016) and included only one maternal family per population. For populations with two maternal families, we arbitrarily selected the one with the lower number, i.e., GAB1, UTC1, NOR511, and SAB1.

### Testing reproductive isolating barriers

To investigate postmating reproductive isolating barriers among species in the *M. tilingii* complex, we performed a crossing experiment using plants from 13 maternal families across 10 populations (maternal families: *M. caespitosa* = 7, *M. minor* = 2, *M. tilingii* = 4; Table S1). For this experiment, we used some of the same individuals as in the morphological and genetic analyses above but supplemented them with full siblings from each maternal family. Intraspecific crosses (C×C, M×M, and T×T, where C = *M. caespitosa,* M = *M. minor*, and T = *M. tilingii*) included two types: 1) crosses within maternal families (i.e., between full sibs), and 2) crosses between maternal families within species. Although we detected some significant differences in postmating isolation between these intraspecific cross types (Table S4), they were likely due to inbreeding depression, as crosses between maternal families usually did better than crosses within maternal families. Therefore, we grouped the two intraspecific cross types for all analyses. Three days prior to each cross, we emasculated maternal parents to avoid contamination from self-pollination. For intraspecific crosses, we generated 62 unique maternal-family cross combinations and 160 total crosses (C×C = 44, M×M = 4, T×T = 14; 1-6 fruits per cross combination). For interspecific crosses, we performed 86 unique and 210 total interspecific crosses (C×M =10, M×C =12, M×T = 8, T×M = 7, T×C = 25, C×T = 24; 1-8 fruits per cross combination; Table S5). We used these crosses to assess the following sequentially-acting postmating reproductive isolating barriers: 1) postmating, prezygotic reproductive isolation, 2) hybrid seed inviability, 3) later-acting hybrid inviability, and 4) hybrid male and female sterility.

#### Postmating, prezygotic isolation

To assess postmating, prezygotic reproductive isolation, we measured seed production per fruit from crosses within and between species. We note that this measure of postmating, prezygotic isolation is likely to be conservative because it reflects only pollen-pistil incompatibilities and not conspecific pollen precedence, which would require mixed pollinations.

We modeled the effect of cross type (i.e., C×C, C×M, M×C, M×M, M×T, T×M, T×T, T×C, C×T) on seed production by fitting a generalized linear model (GLM) with a Gamma distribution using the glm function in the ‘lme4’ package implemented in R (Bates et al. 2007). In this model, the response variable was the number of seeds produced per fruit and the fixed factors were maternal species, paternal species, and their interaction. To determine whether fixed factors and interactions significantly affected the variance of seed production, we computed an ANOVA test using the anova function in the ‘car’ package in R with type III sums of squares, which corrects for unbalanced sample sizes and implements likelihood-ratio chi-square tests for GLMs (Fox et al. 2012). We calculated least-squares means (lsmeans) using the emmeans function in the ‘emmeans’ package in R and performed pairwise comparisons between all cross types (Lenth and Lenth 2018). We used a post-hoc Tukey method adjustment to determine which of the nine cross types differed significantly in the total number of seeds produced.

#### Seed viability

We used two different methods as a proxy for measuring seed viability. First, we performed a visual seed assessment. Recent studies in *Mimulus* have shown that inviable hybrid seeds are often darkened and/or shriveled (Garner et al. 2016, Oneal et al. 2016, Coughlan et al. 2020). Following these studies, we scored round, plump seeds as fully developed and seeds with irregular phenotypes (darkened, shriveled, or wrinkled) as underdeveloped. Second, for a subset of crosses, we also assessed seed viability by scoring seed germination (Table S5). For intraspecific crosses, we measured seed germination rates for 48 unique and 76 total crosses (C×C = 36, M×M = 3, T×T = 9; 1-3 fruits per cross combination). For interspecific crosses, we scored germination for 72 unique and 133 total crosses (C×M = 8, M×C = 7, M×T = 7, T×M = 7, T×C = 20, C×T = 23; 1-4 fruits per cross combination). To determine germination rates, we sowed all seeds from each fruit onto wet paper towels in petri dishes (≤100 seeds per petri dish to avoid overcrowding). Petri dishes were sealed with parafilm, cold-stratified at 4°C for seven days, and then transferred to a growth chamber that provided constant light at 26°C. Ten days later, we scored germination rate as the number of seedlings that had germinated per fruit.

To model the effects of cross type on seed viability, we used generalized linear mixed models (GLMMs). We ran GLMMs for both measures of seed viability (visual assessment and seed germination) with a binomial distribution using the glmer command in the ‘lme4’ package. In this model, we combined the number of viable seeds and the number of inviable seeds into a single response variable using the R function cbind. We set the maternal species, paternal species, and their interaction as fixed factors with their corresponding maternal families set as random factors. Using the anova function with type III sums of squares in R, which applies Wald chi-square tests for mixed models, we computed an ANOVA and determined which fixed factor(s) and interactions significantly contributed to variance of seed viability. We estimated the lsmeans of viable seeds per fruit, performed pairwise comparisons of lsmeans between all cross types, and determined which cross types significantly differed in the number of viable seeds using a post-hoc Tukey method.

#### F1 viability

To investigate later-acting (post-seed) hybrid inviability, we tracked survival to flowering in a subset of the seedlings from the germination tests described in the previous section. We transplanted seedlings from petri dishes into flats with 6-cm cells and transferred them to a 16h, 23°C/8h, 16°C greenhouse. We transplanted 5-16 offspring from each of 27 unique intraspecific crosses (C×C = 17, M×M = 2, T×T = 8; total intraspecific offspring = 315) and 1-23 F1 hybrids from each of 29 unique interspecific crosses (F1s: C×M = 5, M×C = 4, T×M = 4, T×C = 9, C×T = 7; total interspecific offspring = 334). All interspecific cross combinations were represented in these analyses except for M×T, which did not produce viable offspring due to the severe seed inviability phenotype. For each individual, we scored the number of days to flowering as a proxy for viability. For individuals that successfully flowered, we modeled the effect of cross type on days to flower using a GLMM with a Poisson distribution (log link). In this model, we set the response variable as the number of days to flower and the fixed factors as maternal species, paternal species, and their interaction with maternal families treated as random factors. We computed an ANOVA and determined which fixed factor(s) and interactions contributed significantly to variation in days to flowering. We calculated and performed pairwise comparisons of lsmeans and used a post-hoc Tukey test to determine which crosses differed in days to flower. We also visually inspected individuals for signs of necrosis, a plant phenotype that is normally associated with environmental stresses (e.g., pathogen attack) but that can manifest in the absence of pathogens due to hybrid incompatibilities (Bomblies and Weigel 2007).

#### F1 sterility

Finally, using a subset of the intraspecific and hybrid offspring grown to flowering, we investigated both male and female fertility. We assessed male fertility in 4-14 offspring from each of 27 unique intraspecific crosses (C×C = 17, M×M = 2, T×T = 8; total intraspecific offspring = 206) and 4-13 F1 hybrids from each of 28 interspecific crosses (F1s: C×M = 5, M×C = 4, T×M = 4, T×C = 9, C×T = 6; total interspecific offspring = 193). For each individual, we collected anthers from 1-3 of the first four flowers and suspended the pollen in a lacto-phenol aniline blue stain, which stains viable pollen a dark blue color. To estimate pollen viability for each individual, we determined the proportion of viable pollen grains from a haphazard sample of about 100 pollen grains per flower. In a few cases, flowers did not produce functional anthers or pollen (Table S6); these flowers were excluded from further analyses. We modeled whether cross type had a significant effect on pollen viability using a GLMM with a binomial distribution. We combined the number of viable pollen grains and inviable pollen grains into a single variable using the R function cbind and used this as our response variable. Similar to our previous models, we assigned the fixed factors as maternal species, paternal species, and their interaction, with the corresponding maternal families set as random factors. We determined which fixed factors and interactions contributed significantly to variation in pollen viability with ANOVA and estimated pollen viability lsmeans for each cross type. We performed pairwise comparisons of pollen viability lsmeans and determined which cross types differed significantly using a post-hoc Tukey test.

To investigate female fertility, we performed supplemental hand-pollinations on intraspecific and hybrid offspring using one or both of their fertile parents as pollen donors. For each of these hand-pollinations, we counted the number of seeds produced per fruit. We used this approach to assess female fertility in 2-9 offspring from each of 27 unique intraspecific crosses (C×C = 17, M×M = 2, T×T = 8; total intraspecific offspring = 119, 1-3 fruits per individual) and 1-7 F1 hybrids from each of 27 interspecific crosses (F1s: C×M = 5, M×C = 4, T×M = 4, T×C = 9, C×T = 5; total interspecific offspring = 112, 1-4 fruits per individual). To model whether cross type affects F1 seed set, we used a GLMM with a Poisson error distribution (log link). In this model, we first averaged the number of seeds per fruit for each individual, rounded the values to the nearest whole number, and set this as our response variable. The fixed factors of this model were the maternal and paternal species and their interaction, with the maternal families as random factors. We determined the fixed factor(s) and interactions that contributed significantly to F1 seed set variance with ANOVA. Then, we calculated the lsmeans, performed pairwise comparisons of lsmeans, and determined which cross types differed in seed set using a post-hoc Tukey method.

## RESULTS

### Species in the *M. tilingii* complex are morphologically and genetically divergent

To characterize morphological variation within the *M. tilingii* species complex, we grew plants from 16 maternal families together in a common garden. The three putative species within the *M. tilingii* complex showed clear morphological differences in a suite of floral and vegetative traits (Table S2), with a linear discriminant analysis (LDA) separating *M. caespitosa*, *M. minor,* and *M. tilingii* into three non-overlapping clusters (Figure 2, Table S7). Indeed, the three proposed species assignments were identical to the classes predicted by the LDA model (100% of the plants were classified correctly, Table S8).

**Figure 2.**
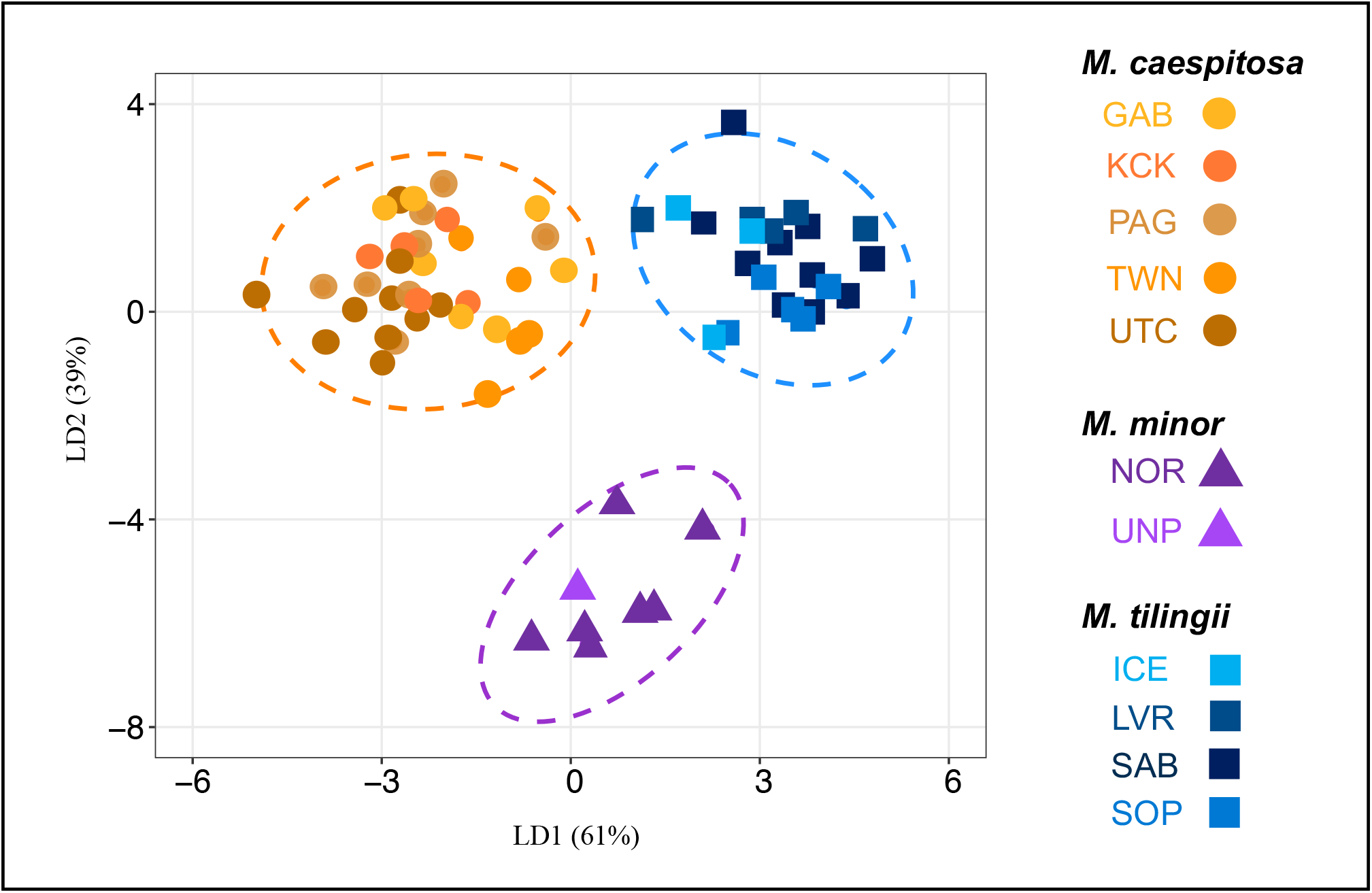
Linear discriminant analysis shows clear morphological differentiation based on floral and vegetative traits measured in a common garden among the three putative species in the *M. tilingii* complex. *M. caespitosa* samples are indicated with circles colored in shades of orange, *M. minor* samples are indicated with triangles colored in shades of purple, and *M. tilingii* samples are indicated with squares colored in shades of blue. Dashed ellipses represent 95% confidence intervals, with corresponding colors.

In addition to these phenotypic differences, patterns of genome-wide variation provide strong support for the existence of three genetically distinct species within the *M. tilingii* complex. A neighbor-joining tree shows the *M. tilingii* complex forms a monophyletic group, which is further separated into three clades corresponding to *M. caespitosa*, *M. minor,* and *M. tilingii* (Figure 3A). Additionally, a principal component analysis reveals genetic structure among species within the *M. tilingii* complex: PC1 (24.30%) splits *M. caespitosa* from *M. minor* and *M. tilingii*, while PC2 (21.89%) separates all three species (Figure 3B). To support these qualitative inferences of genetic structure, we calculated pairwise sequence divergence at fourfold degenerate synonymous sites among *Mimulus* species (Figure 3C, Table S9). Interspecific divergence between *M. caespitosa* and *M. minor* (*d*_*s*_ = 3.78% [3.66%—3.91%]) well exceeds diversity within either species (*M. caespitosa*: π_s_ = 1.2% [1.15%—1.25%]; *M. minor:* π_s_ = 1.04%). Although nucleotide diversity within *M. tilingii* (π_s_ = 3.19% [3.07%—3.31%]) was much higher; interspecific divergence involving this species and *M. caespitosa* (*d*_*s*_ = 4.39% [4.37%—4.41%]) or *M. minor* (*d*_*s*_ = 4.25% [4.23%—4.28%]) was greater still. Using these values and assuming that current levels of diversity within *M. tilingii* approximate levels in the ancestral population, we estimate the species split time (*T*_*S*_) between *M. tilingii* and the other two species to be 377 kya [330ky-425ky] (i.e., following Brandvain et al. 2014: [*T*_*S*_ = *d*_*s* tilxcaes,minor_ – π_s til_]/2μ, where μ = 1.5×10^−8^). Using a similar approach, we estimate 686 kya as the split time between the *M. tilingii* and *M. guttatus* species complexes (approximating ancestral diversity by the average of current diversity in the two complexes; i.e., [*d*_*s* tilxgutt_ – ½ π_s_]/2μ).

**Figure 3.**
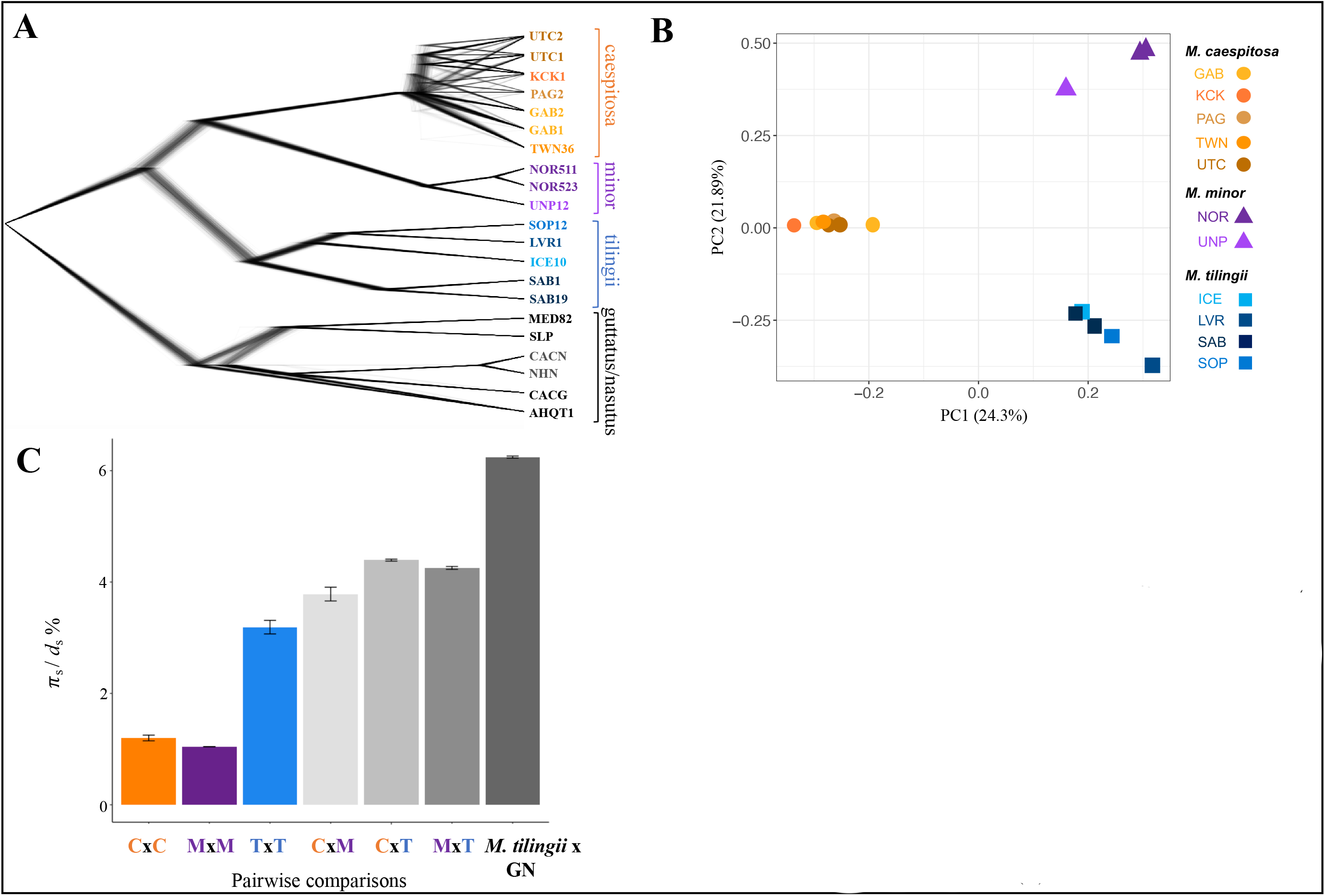
Whole-genome sequence analyses**. A.** Neighbor-joining tree representing the genetic relationships for seven *Mimulus caespitosa*, five *M. tilingii*, three *M. minor,* four *M. guttatus*, and two *M. nasutus* samples, rooted by one *M. dentilobus* sample. The consensus tree was based on 14,000 fourfold degenerate synonymous sites, plotted using a pairwise distance matrix with the nj function in the R package, ape, and smoothed with the function, chronopl, with λ = 1. The distribution of 1000 trees is plotted using the program DensiTree. **B.** Principal component analysis separates species in the *M. tilingii* complex based on genetic relatedness. The PCA uses the same SNP data as the nj tree, but excludes *M. guttatus, M. nasutus, and M. dentilobus. Mimulus caespitosa* samples are shown as round data points colored in shades of orange, *M. minor* samples are shown as triangular data points in shades of purple, and *M. tilingii* samples are shown as squared data points in shades of blue. Note that some populations have two maternal lines. **C.** Average pairwise sequence divergence and +/− SE at fourfold degenerate synonymous sites among *Mimulus* taxa: *M. caespitosa* (C), *M. minor* (M)*, M. tilingii* (T). The darkest gray bar (*M. tilingii* x GN) includes all pairwise sequence comparisons between the three species within the *M. tilingii* complex and *M. guttatus* (G) and *M. nasutus* (N) samples.

### *M. tilingii* species show strong postmating reproductive isolation

To determine the extent of postmating reproductive isolation among the three putative species within the *M. tilingii* complex, we performed a crossing experiment using plants from 10 populations (13 maternal families; *M. caespitosa* = 7, *M. minor* = 2, *M. tilingii* = 4; Table S1). We assessed several sequentially-acting postmating reproductive isolating barriers: 1) postmating, prezygotic reproductive isolation, 2) hybrid seed inviability, 3) later-acting hybrid inviability, and 4) hybrid male and female sterility; results for each are presented below.

Our crosses showed no evidence of postmating, prezygotic reproductive isolation among the three *M. tilingii* species. Indeed, the number of seeds produced by interspecific crosses was just as high as the number produced by intraspecific crosses (Figure 4, Table S10), suggesting that interspecific pollen-pistil incompatibilities do not prevent fertilization among these species. Instead, variation in seed production was driven largely by species of the maternal parent: crosses with *M. minor* as the maternal parent produced 38% more seeds per fruit than crosses with *M. caespitosa* as the maternal parent and 30% more than crosses with *M. tilingii* as the maternal parent (Figure 4, Table S10, Figure S2).

**Figure 4.**
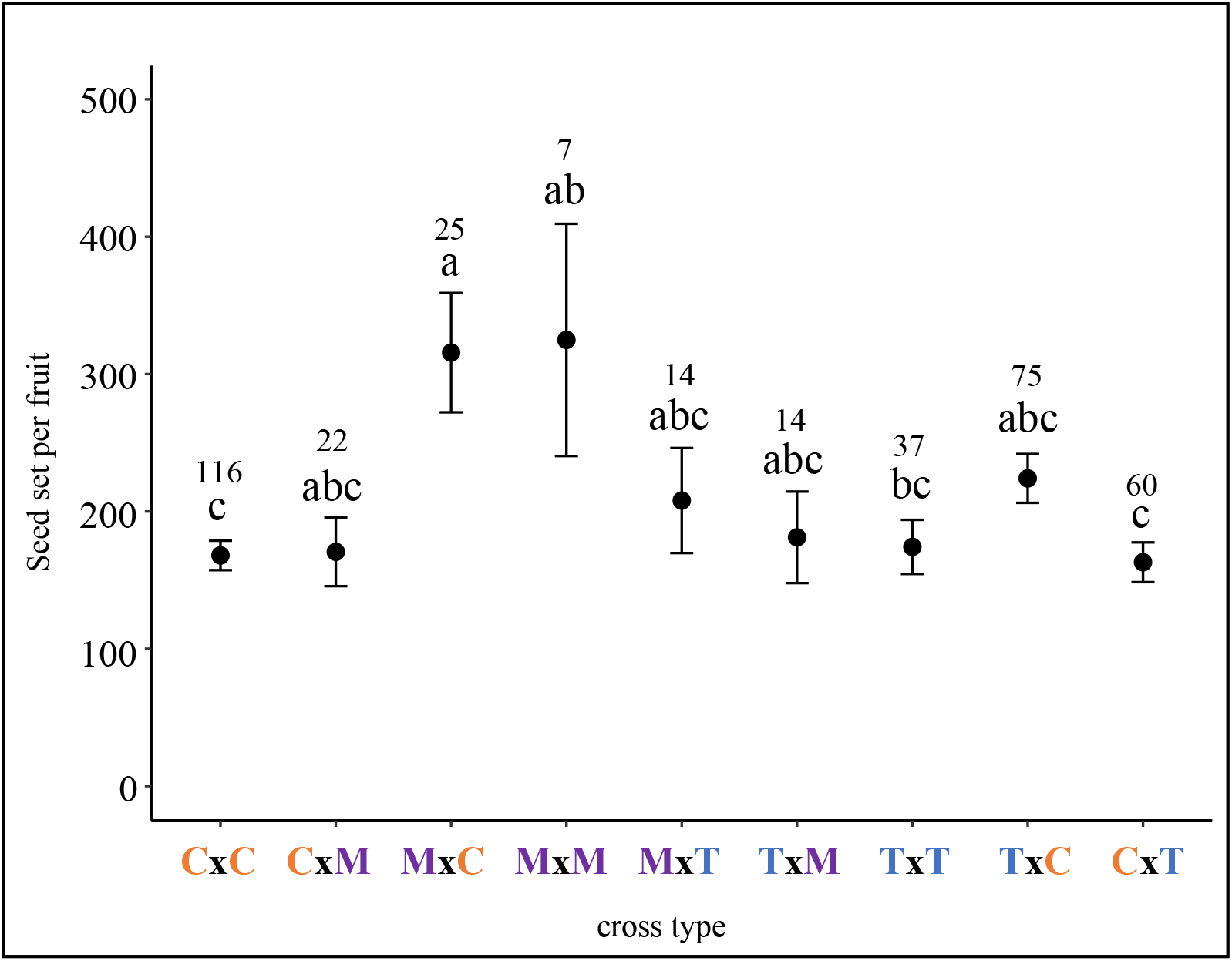
Intraspecific and interspecific seed set per fruit for cross types among *M. caespitosa* (C), *M. minor* (M), and *M. tilingii* (T). The first letter in each cross type indicates the maternal species. Least square means for each cross type are given with +/− SE. Least square means denoted by a different letter indicate significant differences among cross types (*P* <0.05) determined by post-hoc Tukey method. Sample sizes assessed for each cross type are listed above letters.

In contrast to postmating, prezygotic isolation, we discovered very strong hybrid seed inviability in certain crosses within the *M. tilingii* complex using both seed viability measures (visual assessment and germination). In our visual assessment of seed viability, when *M. tilingii* acted as the paternal parent, interspecific crosses produced few to no fully developed seeds per fruit (F1 seed viability: C×T = 19%, M×T < 1%; Figure 5A, Table S10). When the same crosses were performed in the reciprocal direction, the proportion of fully developed hybrid seed was much higher (F1 seed viability: T×C = 89%, T×M = 72%). Hybrid seeds were also mostly fully developed in both reciprocal crosses of *M. caespitosa* and *M. minor* (F1 seed viability: C×M= 96%, M×C = 84%; Figure 5A). Variation in germination rates among cross types largely mirrored patterns of visually assessed seeds (i.e., the rank order among cross types did not change; Figure 5B, Table S12). In sum, hybrid seed inviability is a strong reproductive isolating barrier in one crossing direction between *M. tilingii* and *M. caespitosa* or *M. minor*.

**Figure 5.**
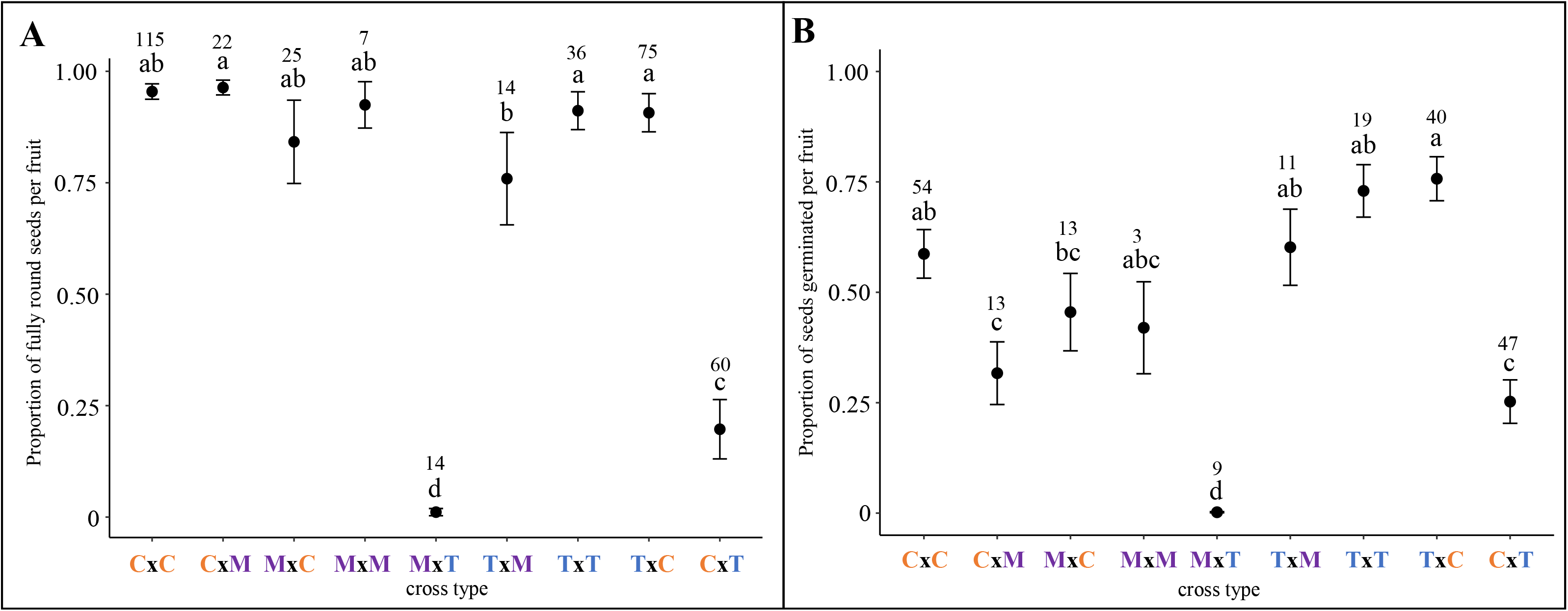
Intraspecific and interspecific seed viability for crosses among *M. caespitosa* (C), *M. minor* (M), and *M. tilingii* (T). The first letter in each cross type indicates the maternal species. Least square means for cross types are given with +/− SE. Least square means denoted by a different letter indicate significant differences among cross types (*P* <0.05) determined by post-hoc Tukey method. Sample sizes assessed for each cross type are listed above letters. **A.** Proportion of fully developed seeds per fruit (visual assessment). **B.** Proportion of seeds that germinated per fruit.

Next, we assessed the viability of hybrids that survived to the seedling stage. Once established as seedlings, all progeny of intraspecific crosses, and most hybrid progeny of interspecific crosses, survived to flowering (Table S6). We detected no evidence of F1 hybrid inviability between *M. caespitosa* and *M. minor*: 100% of C×M and M×C F1 hybrids produced flowers (*N* = 57 and 56, respectively). In fact, C×M F1 hybrids flower much earlier (~13 days) than progeny of *M. caespitosa* crosses (Table S13). Similarly, 100% of F1 hybrids between *M. tilingii* and *M. minor* flowered and showed no delay in flowering time relative to progeny of intraspecific crosses (*N* = 54 for T×M; severe hybrid seed inviability precluded generating F1 hybrids in the reciprocal direction, Table S13). However, one class of F1 hybrids – those produced from crosses between *M. caespitosa* and *M. tilingii* – did show evidence of inviability: 18% of C×T and T×C F1 hybrids did not survive to flowering because they were severely necrotic (*N* = 60 and 107, respectively, Figure S2; Table S6). It is important to note that this F1 hybrid necrosis phenotype was not segregating in all *M. caespitosa-M. tilingii* crosses. Instead, the 18% frequency is due to a high proportion of necrotic F1 hybrids between particular maternal families of *M. caespitosa* and *M. tilingii* (i.e., 25-100% F1 necrosis in crosses between *M. caespitosa* GAB1 or UTC1 and *M. tilingii* ICE10; Table S6). Thus, although hybrid inviability is not fixed between species of the *M. tilingii* complex, it can be a strong postzygotic isolating barrier in certain interspecific crosses.

Finally, for hybrids and intraspecific progeny that survived to flowering, we examined both male and female fertility. Strikingly, we discovered strong male sterility in both of the reciprocal F1 hybrids from all three interspecific crosses: pollen viability was much lower in all F1 hybrids than in the progeny of intraspecific crosses (Figure 6A, Table S14). Male sterility was particularly severe in F1 hybrids with *M. minor* as a parent (C×M, M×C, and T×M), which showed a 96% reduction in pollen viability compared to intraspecific crosses. Female sterility was also remarkably strong in all tested F1 hybrids (Figure 6B; Table S15). Reciprocal F1 hybrids between *M. caespitosa* and *M. tilingii* produced 81% fewer seeds per fruit than parental intraspecific crosses (T×T and C×C). As with male sterility, F1 hybrids with *M. minor* as a parent showed particularly severe female sterility, with a 99% reduction in F1 seed set compared to intraspecific crosses. Taken together, these results indicate that F1 sterility – through both male and female functions – is an extremely strong postzygotic isolating barrier between species in the *M. tilingii* complex.

**Figure 6.**
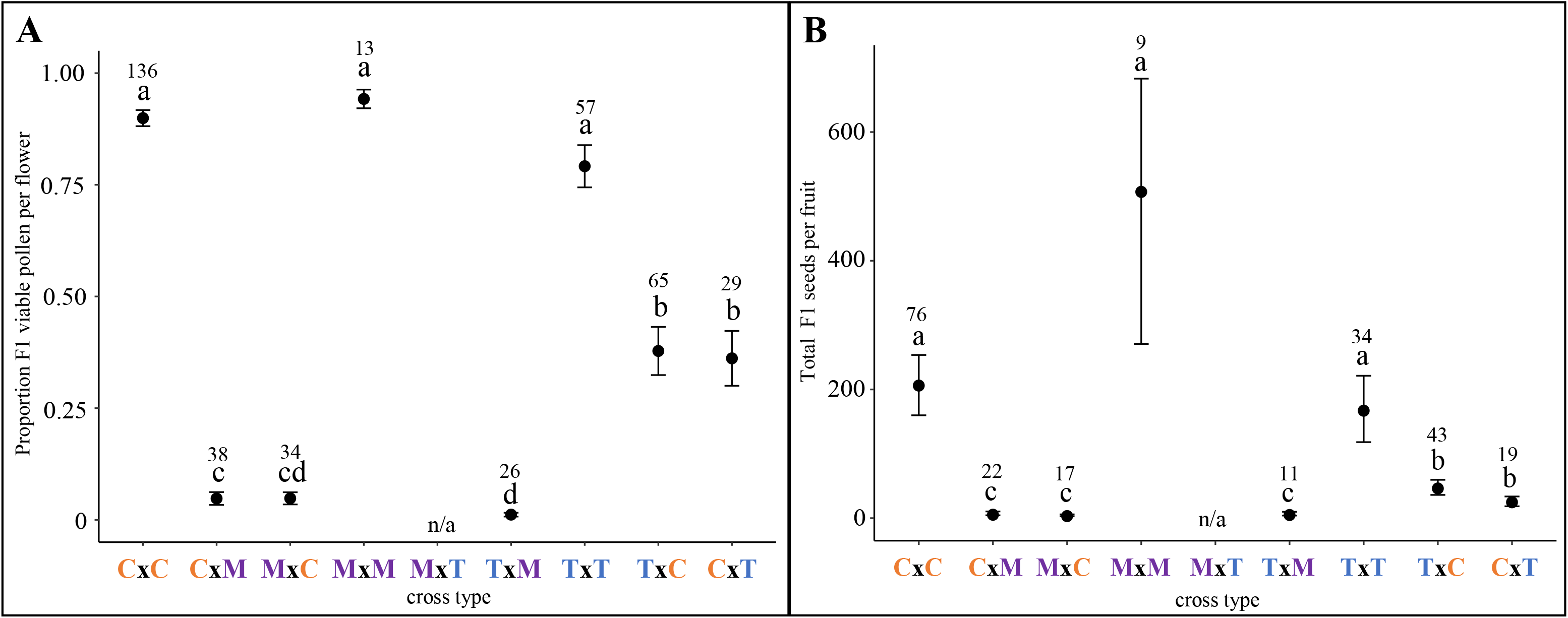
F1 intraspecific and interspecific fertility for crosses among *M. caespitosa* (C), *M. minor* (M), and *M. tilingii* (T). The first letter in each cross type indicates the maternal species. Least square means for each cross type are given with +/− SE. Least square means denoted by a different letter indicate significant differences among cross types (*P* <0.05) determined by post-hoc Tukey method. Sample sizes assessed for each cross type are listed above letters. There is no data for the M x T cross type due to the severe seed lethality phenotype. **A.** Proportion of F1 pollen viability per flower. **B.** Total F1 seeds produced per fruit.

## DISCUSSION

A fundamental goal in evolutionary biology is understanding how new species evolve. In this study, we determined that closely related yet morphologically distinct species within the *M. tilingii* complex (*M. caespitosa*, *M. minor*, and *M. tilingii*) genetically diverged approximately 377kya. Additionally, we discovered that a cross between any species pair within the *M. tilingii* complex results in near complete reproductive isolation by several postzygotic barriers, including hybrid seed inviability, hybrid necrosis, and hybrid male and female sterility. Below, we discuss the possibility that strict allopatry among these montane species within the *M. tilingii* complex might have facilitated the evolution of this strikingly high number of hybrid incompatibilities.

In this study, the first severe postzygotic barrier we found between certain species within the *M. tilingii* complex was hybrid seed inviability. In flowering plants, hybrid seed inviability is a common feature of interploidy and interspecific crosses (Scott et al. 1998, Rebernig et al. 2015, Roth et al. 2018) and, in fact, has evolved multiple times across the *Mimulus* genus (Vickery 1978, Garner et al. 2016, Oneal et al. 2016, Coughlan et al. 2020). Often, hybrid seed inviability is caused by a defective endosperm—a tissue critical for transferring maternal nutrients to the developing embryo (Köhler et al. 2010, Lafon-Placette and Köhler 2016, Brink and Cooper 1947). The endosperm also serves as the primary tissue of genomic imprinting, which is parent-of-origin dependent gene expression due to differential epigenetic modifications established during male and female gametogenesis (Köhler et al. 2012). Classic theory suggests that the evolution of imprinted genes might be driven by parental conflict over maternal investment in the endosperm (Haig and Westoby 1989). In principle, misregulation of imprinted genes provides a mechanistic explanation for the common observation that the seeds of reciprocal interspecific crosses often show phenotypic differences (Haig and Westoby 1991). In like manner, we show parent-of-origin effects on hybrid seed inviability among species in the *M. tilingii* complex; notably, seeds are mostly inviable when *M. tilingii* acts as the pollen donor in any interspecific cross. Reciprocal differences in seed viability are a hallmark of endosperm defects (Haig and Westoby 1991), and although we do not show a defective endosperm as a mechanistic cause, patterns of seed viability among *M. tilingii* species (Figure 5) and preliminary developmental work (Sandstedt and Sweigart, unpublished) suggests the endosperm is involved.

Crosses between species with divergent mating systems (i.e., self-fertilizers and outcrossers) can result in reciprocal seed phenotypes, which might be driven by differences in strength of conflict (weak inbreeder/strong outbreeder [WISO] hypothesis; Brandvain and Haig 2005). In the case of the *M. tilingii* species complex, a shift towards selfing in *M. minor* and *M. caespitosa* could explain reciprocal differences in seed viability in hybrid crosses with *M. tilingii*. Apart from mating system differences, the strength of parental conflict may depend on other factors that influence genetic variation, including demographic history or vegetative propagation (i.e., stolons) that can lead to clonal reproduction. Moreover, the genetic and evolutionary basis of hybrid seed inviability among *M. tilingii* species might be much more complex. We note that although hybrid seed inviability appears to be mostly species-wide, one *M. caespitosa* maternal line (GAB1) consistently produced viable seeds when crossed reciprocally with *M. tilingii*, suggesting that causal genetic loci may be polymorphic within species.

In addition to early-acting hybrid seed inviability between species within the *M. tilingii* complex, we found later-acting inviability in the form of hybrid necrosis. This plant syndrome is associated with a suite of phenotypes including cell death, wilting, yellowing, chlorosis, reduced growth rates, and often lethality (Bomblies and Weigel 2007). In crosses between *M. tilingii* and *M. caespitosa*, we discovered severe F1 hybrid necrosis: plants produced unusually small buds that failed to develop into flowers, followed by plant senescence (observed; Figure S3). Hybrid lethality can readily evolve in many plant systems and has been reported several times in *Mimulus* (Macnair and Christie 1983, Lowry et al. 2008, Wright et al. 2013, Zuellig and Sweigart 2018). As in other plant taxa (Sicard et al. 2015, Zuellig and Sweigart 2018, Macnair and Christie 1983), we observed variation in the genetic basis of hybrid lethality within *M. tilingii* species— only specific maternal lines in combination give rise to hybrid lethal offspring (i.e., UTC1 and GAB1 in combination with ICE10). Often, hybrid necrosis is caused when incompatible disease resistance genes (i.e., R genes) against bacterial or fungal pathogens facilitate an autoimmune response (Chae et al. 2016). Disease resistance genes are thought to evolve rapidly in response to pathogen pressure; they exhibit exceptional variation in nucleotide sequence, high copy number, and gene expression (Jacob et al. 2013). Additionally, in natural plant populations, disease resistance genes often show signatures of balancing selection and diversifying selection (Karasov et al. 2014), which might explain why causal genetic loci are polymorphic within *M. tilingii* and *M. caespitosa* species. Future experiments are needed to determine the molecular genetic basis of hybrid necrosis in the *M. tilingii* complex and whether divergence of disease resistance genes underlies this barrier.

Lastly, we show that viable F1 hybrids generated from crosses among species within the *M. tilingii* complex are severely male and female sterile, especially when *M. minor* is involved. Hybrid sterility is a common reproductive barrier across plants and animals and its genetic basis can vary from simple to complex (Kubo et al. 2008, Sweigart et al. 2006, Lai et al. 2005). Many factors have been implicated as causes underlying hybrid male sterility, including cytonuclear incompatibilities, chromosomal rearrangements, interactions among nuclear genes, or a combination of these factors (Bomblies 2010). For example, in closely related sunflower species, severe hybrid sterility was genetically mapped to karyotypic differences between species as well as genic interactions in non-rearranged regions (Lai et al. 2005). Because both male and female hybrid sterility are strong among species within the *M. tilingii* complex, we speculate that chromosomal rearrangements and/or multiple independent Dobzhansky-Muller incompatibilities may underlie these barriers. Although we cannot completely rule out slight variation in chromosome number as a potential cause for hybrid incompatibilities among species within the *M. tilingii* complex, preliminary chromosome squashes suggest no differences in ploidy (n=14; data not shown).

What factors might explain the evolution of multiple strong reproductive barriers among these closely related members of the *M. tilingii* species complex? During and following Pleistocene glaciation, it is possible that gene flow was severely limited among species in the *M. tilingii* complex that were confined to distinct mountain ranges, facilitating the accumulation of hybrid incompatibilities and other genetic differences. Although the *M. tilingii* complex is as genetically variable as the closely related and well-studied *M. guttatus* species complex (Brandvain et al. 2014), it shows much stronger F1 postzygotic isolation. Many hybrid incompatibilities have been identified within and between members of the *M. guttatus* complex, including several that affect F2 hybrids or backcross hybrids (hybrid lethality in Zuellig and Sweigart 2018, hybrid sterility in Sweigart et al. 2006, Fishman and Willis 2006) though some species pairs give rise to F1 hybrid seed inviability and various levels of hybrid lethality (Gardner and Macnair 2000, Macnair and Christie 1983, Wright et al. 2013). In addition, species in the *M. guttatus* complex have overlapping distributions throughout most of Western North America, and there is evidence for substantial introgression in regions of sympatry (Brandvain et al. 2014, Kenney and Sweigart 2016, Zuellig and Sweigart 2018). When interspecific gene flow is present, theory suggests that neutrally evolving hybrid incompatibility alleles may be purged from species because their deleterious effects become exposed in hybrids (Gavrilets 1997, Kondrashov 2003, Bank et al. 2012, Muir and Hahn 2015). Perhaps, then, extensive gene flow between species in the *M. guttatus* complex explains its lower prevalence of F1 postzygotic barriers, and strict allopatry in the *M. tilingii* complex might explain why much stronger postzygotic isolation has evolved among its species. In plants, closely related species often show extensive range overlap (Baack et al. 2015), yet it is unclear how such overlap will impact the strength of intrinsic postzygotic isolation. More studies are needed to explicitly test the effect of interspecific gene flow on the relative strength of prezygotic vs. postzygotic barriers in young species pairs.

Along with these strong intrinsic F1 postzygotic barriers, it is entirely possible that prezygotic and extrinsic postzygotic barriers might also have evolved among allopatric species within the *M. tilingii* complex. For example, even though we did not find evidence of postmating, prezygotic isolation in our study, we did not test for conspecific pollen precedence, which has been shown to partially isolate other closely related *Mimulus* species (Diaz and Macnair 1999; Ramsey et al. 2003, Fishman et al. 2008). Additionally, we find that patterns of morphological and genetic variation among *M. tilingii* species might be driven, at least in part, by mating system divergence. Shifts in mating system are common across the *Mimulus* genus and other flowering plants and can act as a strong premating barrier (Rieseberg & Willis 2007). Although members of the *M. tilingii* species complex appear to be predominantly outcrossing, the rate of selfing within and between species varies and can be as high as 30% (Ritland 1989; Ritland and Ritland 1989). Compared to *M. tilingii*, individuals from both *M. minor* and *M. caespitosa* show a relative decrease in anther-stigma distance and corolla width (Table S2), two traits that can promote selfing via contact between the stigma and anthers. Consistent with a transition toward increased selfing in these species, nucleotide diversity between populations of *M. caespitosa* and *M. minor* was only a third that of *M. tilingii* (Figure 3C). Additionally, nucleotide diversity within the NOR population of *M. minor* was 0.02% (*N* = 2), which represents a 50-fold reduction in intrapopulation variation compared to *M. caespitosa* and *M. tilingii* (Table S9). Although these results might suggest an increased propensity for selfing in *M. minor*, we note that one maternal line belonging to *M. minor* had the highest individual heterozygosity compared to all other sequenced lines in this study (UNP12; Table S1). Further, some patterns may be explained by the fact that we have only two *M.* minor populations and a narrower sampling distribution for both *M. caespitosa* and *M. minor*.

Although it is tempting to speculate that speciation in the *M. tilingii* species complex has been driven in large part by postzygotic reproductive isolation, more work will be needed to understand the evolutionary causes and consequences of F1 postzygotic barriers in nature. The exact geographical distributions of members in the *M. tilingii* species complex are not well defined and we do not yet know whether these species occasionally come into secondary contact. Additionally, although we know species within the *M. tilingii* complex are restricted to high elevations, more investigation is needed to determine whether these montane environments are ecologically distinct and whether species within the *M. tilingii* complex have evolved premating barriers associated with divergent adaptation. In conclusion, species in the *M. tilingii* complex are closely related, yet genetically and morphologically distinct. Notably, this system is rich with possibilities to investigate the genetics and evolution of reproductive isolation in montane, allopatric species early in divergence.

## Author Contributions

Research conceived and designed by G.D.S. and A.L.S., data collected and analyzed by G.D.S., manuscript written by G.D.S. and A.L.S., and C.A.W. made original collections for plants used in this work and made comments on the manuscript.

## Acknowledgments

The Rocky Mountain Biological Laboratory provided permission for sample collections in Colorado State. We thank Mason Lin, Sydney Dilworth, and Tylanna Baker for data collection and Taylor Harrell for plant illustrations (Figure S1). We are grateful to Jill Anderson, Wolfgang Lukowitz, David Hall, Robert Schmitz, Robert Franks, Rachel Kerwin, Samuel Mantel, and Makenzie Whitener for helpful discussions. David Hall, Robert Franks, Samuel Mantel, Matt Farnitano, and Makenzie Whitener provided valuable comments and improved the quality of the manuscript. This work was supported by the National Institutes of Health T32 Fellowship [GM007103] to G.D.S., the Jan and Kirby Alton Fellowship [Department of Genetics, UGA] to G.D.S., and National Science Foundation grants [DEB-1350935 and DEB-1856180] to A.L.S.

The authors declare no competing interests.

## Data Accessibility Statement

Data will be made available on Dryad Digital Repository and whole genome sequence data will be uploaded to NCBI Short Read Archive.

## REFERENCES

Andrews S. 2010. FastQC: a quality control tool for high throughput sequence data. Available online at: http://www.bioinformatics.babraham.ac.uk/projects/fastqc

Baack, E., Melo, M. C., Rieseberg, L. H., & Ortiz-Barrientos, D. 2015. The origins of reproductive isolation in plants. New Phytologist, 207(4), 968–984.

Bank, C., Hermisson, J., & Kirkpatrick, M. 2012. Can reinforcement complete speciation? Evolution: International Journal of Organic Evolution, 66(1), 229–239.

Bates, D., Sarkar, D., Bates, M. D., & Matrix, L. 2007. The lme4 package. R package version, 2(1), 74.

Bolger, A. M., Lohse, M., & Usadel, B. 2014. Trimmomatic: a flexible trimmer for Illumina sequence data. Bioinformatics, 30(15), 2114–2120.

Bomblies, K. 2010. Doomed lovers: mechanisms of isolation and incompatibility in plants. Annual review of plant biology, 61, 109–124.

Bomblies, K., & Weigel, D. 2007. Hybrid necrosis: autoimmunity as a potential gene-flow barrier in plant species. Nature Reviews Genetics, 8(5), 382–393.

Bouckaert, R. R. 2010. DensiTree: making sense of sets of phylogenetic trees. Bioinformatics, 26(10), 1372–1373.

Brandvain, Y., & Haig, D. 2005. Divergent mating systems and parental conflict as a barrier to hybridization in flowering plants. The American Naturalist, 166(3), 330–338.

Brandvain, Y., Kenney, A. M., Flagel, L., Coop, G., & Sweigart, A. L. 2014. Speciation and introgression between *Mimulus* nasutus and *Mimulus guttatus*. PLoS genetics, 10(6), e1004410.

Brink, R. A., & Cooper, D. C. 1947. The endosperm in seed development. The Botanical Review, 13(9), 479–541.

Butlin, R. 1989. Reinforcement of premating isolation. Pp. 158–179 in Otte, D. and J. A. Endler (eds) Speciation and its consequences. Sinauer Associates, Inc., Sunderland, MA.

Chae, E., Tran, D. T., & Weigel, D. 2016. Cooperation and conflict in the plant immune system. PLoS pathogens, 12(3).

Christie, K., & Strauss, S. Y. 2019. Reproductive isolation and the maintenance of species boundaries in two serpentine endemic Jewelflowers. Evolution, 73(7), 1375–1391.

Coughlan, J. M., Brown, M. W., & Willis, J. H. 2020. Patterns of Hybrid Seed Inviability in the *Mimulus guttatus* sp. Complex Reveal a Potential Role of Parental Conflict in Reproductive Isolation. Current Biology, 30(1), 83–93.

Coyne, J. A., and H. A. Orr. 1989. Patterns of speciation in *Drosophila*. Evolution 43:362–381.

Coyne, J.A., and H. A. Orr. 2004. Speciation. Sinauer Associates, Sunderland, MA.

Darwin, C. R. 1859. The Origin of species. 6th ed. John Murray, London.

Diaz, A., & Macnair, M. R. 1999. Pollen tube competition as a mechanism of prezygotic reproductive isolation between *Mimulus* nasutus and its presumed progenitor *M. guttatus*. The New Phytologist, 144(3), 471–478.

Dobzhansky, T. 1937. Genetic nature of species differences. The American Naturalist, 71(735), 404–420.

Dobzhansky, T. 1951. Genetics and the origin of species. 3rd ed. Columbia Univ. Press, New York

Doyle, J. J., & Doyle, J. L. 1987. CTAB DNA extraction in plants. Phytochemical Bulletin, 19, 11–15.

Fishman, L., Aagaard, J., & Tuthill, J. C. 2008. Toward the evolutionary genomics of gametophytic divergence: patterns of transmission ratio distortion in monkeyflower (*Mimulus*) hybrids reveal a complex genetic basis for conspecific pollen precedence. Evolution: International Journal of Organic Evolution, 62(12), 2958–2970.

Fishman, L., & Sweigart, A. L. 2018. When two rights make a wrong: the evolutionary genetics of plant hybrid incompatibilities. Annual review of plant biology, 69, 707–731.

Fishman, L., & Willis, J. H. 2006. A cytonuclear incompatibility causes anther sterility in *Mimulus* hybrids. Evolution, 60(7), 1372–1381.

Fox, J., Weisberg, S., Adler, D., Bates, D., Baud-Bovy, G., Ellison, S., Firth, D., Friendly, M., Gorjanc, G., Graves, S. and Heiberger, R. 2012. Package ‘car’. Vienna: R Foundation for Statistical Computing

Gardner, M., & Macnair, M. 2000. Factors affecting the co-existence of the serpentine endemic *Mimulus nudatus* Curran and its presumed progenitor, *Mimulus guttatus* Fischer ex DC. Biological Journal of the Linnean Society, 69(4), 443–459.

Garner, A. G., Kenney, A. M., Fishman, L., & Sweigart, A. L. 2016. Genetic loci with parent of-origin effects cause hybrid seed lethality in crosses between *Mimulus* species. New Phytologist, 211(1), 319–331.

Gavrilets, S. 1997. Hybrid zones with Dobzhansky-type epistatic selection. Evolution, 51(4), 1027–1035.

Grant, A.L. 1924. A monograph of the genus *Mimulus*. Ann. Missouri Bot. Gard. 11:99–389

Haig, D., & Westoby, M. 1989. Parent-specific gene expression and the triploid endosperm. The American Naturalist, 134(1), 147–155.

Haig, D., & Westoby, M. 1991. Genomic imprinting in endosperm: its effect on seed development in crosses between species, and between different ploidies of the same species, and its implications for the evolution of apomixis. Philosophical Transactions: Biological Sciences, 1–13.

Hopkins, R. 2013. Reinforcement in plants. New Phytologist, 197(4), 1095–1103.

Ishizaki, S., Abe, T., & Ohara, M. 2013. Mechanisms of reproductive isolation of interspecific hybridization between *Trillium camschatcense* and *T. tschonoskii* (Melanthiaceae). Plant Species Biology, 28(3), 204–214.

Jacob, F., Vernaldi, S., & Maekawa, T. 2013. Evolution and conservation of plant NLR functions. Frontiers in immunology, 4, 297.

Karasov, T. L., Horton, M. W., & Bergelson, J. 2014. Genomic variability as a driver of plant-pathogen coevolution? Current opinion in plant biology, 18, 24–30.

Kenney, A. M., & Sweigart, A. L. 2016. Reproductive isolation and introgression between sympatric *Mimulus* species. Molecular ecology, 25(11), 2499–2517.

Köhler, C., Scheid, O. M., & Erilova, A. 2010. The impact of the triploid block on the origin and evolution of polyploid plants. Trends in Genetics, 26(3), 142–148.

Köhler, C., Wolff, P., & Spillane, C. 2012. Epigenetic mechanisms underlying genomic imprinting in plants. Annual review of plant biology, 63, 331–352.

Kondrashov, A. S. 2003. Accumulation of Dobzhansky-Muller incompatibilities within a spatially structured population. Evolution, 57(1), 151–153.

Kubo, T., Yamagata, Y., Eguchi, M., & Yoshimura, A. 2008. A novel epistatic interaction at two loci causing hybrid male sterility in an inter-subspecific cross of rice (*Oryza sativa* L.). Genes & genetic systems, 83(6), 443–453.

Lafon-Placette, C., & Köhler, C. 2016. Endosperm-based postzygotic hybridization barriers: developmental mechanisms and evolutionary drivers. Molecular Ecology, 25(11), 2620–2629.

Lai, Z., Nakazato, T., Salmaso, M., Burke, J. M., Tang, S., Knapp, S. J., & Rieseberg, L. H. 2005. Extensive chromosomal repatterning and the evolution of sterility barriers in hybrid sunflower species. Genetics, 171(1), 291–303.

Lenth, R., & Lenth, M. R. 2018. Package ‘lsmeans’. The American Statistician, 34(4), 216–221.

Li, H., and Durbin, R. 2009. Fast and accurate short read alignment with Burrows-Wheeler transform. Bioinformatics 25:1754–1760.

Li, H., Handsaker, B., Wysoker, A., Fennell, T., Ruan, J., Homer, N., Marth, G., Abecasis, G & Durbin, R. 2009. The sequence alignment/map format and SAMtools. Bioinformatics, 25(16), 2078–2079.

Li, H. 2013. Aligning sequence reads, clone sequences and assembly contigs with BWA-MEM. arXiv 1303:3997v2.

Lowry, D. B., Modliszewski, J. L., Wright, K. M., Wu, C. A., & Willis, J. H. 2008. The strength and genetic basis of reproductive isolating barriers in flowering plants. Philosophical Transactions of the Royal Society B: Biological Sciences, 363(1506), 3009–3021.

Lowry, D. B., Rockwood, R. C., & Willis, J. H. 2008. Ecological reproductive isolation of coast and inland races of *Mimulus guttatus*. Evolution: International Journal of Organic Evolution, 62(9), 2196–2214.

Lowry, D.B., Sobel, J.M., Angert, A.L., Ashman, T.L., Baker, R.L., Blackman, B.K., Brandvain, Y., Byers, K.J., Cooley, A.M., Coughlan, J.M. and Dudash, M.R. 2019. The case for the continued use of the genus name *Mimulus* for all monkeyflowers. Taxon, 68(4), pp.617–623.

Macnair, M. R., & Christie, P. 1983. Reproductive isolation as a pleiotropic effect of copper tolerance in *Mimulus guttatus*? Heredity, 50(3), 295–302.

Mayr, E. 1942. Systematics and the origin of species. Columbia Univ. Press, New York.

McKenna, A., Hanna, M., Banks, E., Sivachenko, A., Cibulskis, K., Kernytsky, A., Garimella, K., Altshuler, D., Gabriel, S., Daly, M. & DePristo, M. A. 2010. The Genome Analysis Toolkit: a MapReduce framework for analyzing next-generation DNA sequencing data. Genome research, 20(9), 1297–1303.

Morjan, C. L., & Rieseberg, L. H. 2004. How species evolve collectively: implications of gene flow and selection for the spread of advantageous alleles. Molecular ecology, 13(6), 1341–1356.

Muir, C. D., & Hahn, M. W. 2015. The limited contribution of reciprocal gene loss to increased speciation rates following whole-genome duplication. The American Naturalist, 185(1), 70–86.

Mukherjee, B. B., & Vickery, R. K. 1959. Chromosome counts in the section Simiolus of the genus *Mimulus* (Scrophulariaceae). III. Madroño, 15(2), 57–62.

Mukherjee, B. B., & Vickery, R. K. 1960. Chromosome counts in the section Simiolus of the genus *Mimulus* (Scrophulariaceae). IV. Madroño, 15(8), 239–245.

Mukherjee, B. B., & Vickery, R. K. 1962. Chromosome counts in the section Simiolus of the genus *Mimulus* (Scrophulariaceae). V. The chromosomal homologies of *M. guttatus* and its allied species and varieties. Madroño, 16(5), 141–155.

Muller H. J. 1942. Isolating mechanisms, evolution, and temperature. Biological Symposium 6: 71–125.

Nesom, G. L. 2012. Taxonomy of *Erythranthe* sect. Simiola (Phrymaceae) in the USA and Mexico. Phytoneuron, 40, 1–123.

Nesom, G.L. 2013. New distribution records for *Erythranthe* (Phrymaceae). Phytoneuron 2013, 67: 1–15.

Nesom, G.L. 2014. Updated classification and hypothetical phylogeny of *Erythranthe* sect. Simiola (Phrymaceae). Phytoneuron, 2014, 1–6.

Nesom, G.L. 2019. Taxonomic status of *Erythranthe minor* (Phrymaceae). Phytoneuron 2019, 32: 1–7.

Nesom, G. L., Fraga, N. S., Barker, W. R., Beardsley, P. M., Tank, D. C., Baldwin, B. G., & Olmstead, R. G. 2019. Response to “The case for the continued use of the genus name *Mimulus* for all monkeyflowers”.

Noor, M. A. 1999. Reinforcement and other consequences of sympatry. Heredity, 83(5), 503–508.

Okonechnikov, K., Conesa, A., & García-Alcalde, F. 2015. Qualimap 2: advanced multi-sample quality control for high-throughput sequencing data. Bioinformatics, 32(2), 292–294.

Oneal, E., Willis, J. H., & Franks, R. G. 2016. Disruption of endosperm development is a major cause of hybrid seed inviability between *Mimulus guttatus* and *Mimulus* nudatus. New Phytologist, 210(3), 1107–1120.

Ostevik, K. L., Andrew, R. L., Otto, S. P., & Rieseberg, L. H. 2016. Multiple reproductive barriers separate recently diverged sunflower ecotypes. Evolution, 70(10), 2322–2335.

Paradis, E., Claude, J., & Strimmer, K. 2004. APE: analyses of phylogenetics and evolution in R language. Bioinformatics, 20(2), 289–290.

Pennell, F. W. 1951. Mimulus. Illustrated flora of the Pacific states, 3, 688–731.

Ramsey, J., Bradshaw Jr, H. D., & Schemske, D. W. 2003. Components of reproductive isolation between the monkeyflowers *Mimulus lewisii* and *M. cardinalis* (Phrymaceae). Evolution, 57(7), 1520–1534.

Rasband, W.S. 1997. ImageJ, U. S. National Institutes of Health, Bethesda, Maryland, USA, https://imagej.nih.gov/ij/.

Rebernig, C. A., Lafon-Placette, C., Hatorangan, M. R., Slotte, T., & Köhler, C. 2015. Non-reciprocal interspecies hybridization barriers in the *Capsella* genus are established in the endosperm. PLoS genetics, 11(6).

Rieseberg, L. H. 2001. Chromosomal rearrangements and speciation. Trends in ecology & evolution, 16(7), 351–358.

Rieseberg, L. H., & Willis, J. H. 2007. Plant speciation. Science, 317(5840), 910–914.

Ritland, C. and Ritland, K., 1989. Variation of sex allocation among eight taxa of the *Mimulus guttatus* species complex (Scrophulariaceae). American Journal of Botany, 76(12), pp.1731–1739.

Ritland, K. 1989. Genetic differentiation, diversity, and inbreeding in the mountain monkeyflower (*Mimulus* caespitosus) of the Washington Cascades. Canadian Journal of Botany, 67(7), 2017–2024.

Roth, M., Florez-Rueda, A. M., Griesser, S., Paris, M., & Städler, T. 2018. Incidence and developmental timing of endosperm failure in post-zygotic isolation between wild tomato lineages. Annals of botany, 121(1), 107–118.

Schliep, K. P. 2010. phangorn: phylogenetic analysis in R. Bioinformatics, 27(4), 592–593.

Schluter, D. 2001. Ecology and the origin of species. Trends in ecology & evolution, 16(7), 372–380.

Scott, R. J., Spielman, M., Bailey, J., & Dickinson, H. G. 1998. Parent-of-origin effects on seed development in Arabidopsis thaliana. Development, 125(17), 3329–3341.

Sicard, A., Kappel, C., Josephs, E. B., Lee, Y. W., Marona, C., Stinchcombe, J. R., Wright, S.I. & Lenhard, M. 2015. Divergent sorting of a balanced ancestral polymorphism underlies the establishment of gene-flow barriers in *Capsella*. Nature communications, 6, 7960.

Sobel, J. M., Chen, G. F., Watt, L. R., & Schemske, D. W. 2010. The biology of speciation. Evolution: International Journal of organic evolution, 64(2), 295–315.

Stebbins, G. L. 1958. The inviability, weakness and sterility of interspecific hybrids. Adv. Genet. 9:147–215.

Suni, S. S., & Hopkins, R. 2018. The relationship between postmating reproductive isolation and reinforcement in Phlox. Evolution, 72(7), 1387–1398.

Sweigart, A. L., Fishman, L., & Willis, J. H. 2006. A simple genetic incompatibility causes hybrid male sterility in *Mimulus*. Genetics, 172(4), 2465–2479.

Vickery, R.K., Jr. 1974. Crossing barriers in the yellow monkey flowers in the genus *Mimulus* (Scrophulariaceae). Genet. Lect. 3: 33–82.

Vickery, R.K., Jr. 1978. Case studies in the evolution of species complexes in *Mimulus*. Evolutionary Biology. 11: 405–507.

Widmer, A., Lexer, C., & Cozzolino, S. 2009. Evolution of reproductive isolation in plants. Heredity, 102(1), 31–38.

Wright, K. M., Lloyd, D., Lowry, D. B., Macnair, M. R., & Willis, J. H. 2013. Indirect evolution of hybrid lethality due to linkage with selected locus in *Mimulus guttatus*. PLoS biology, 11(2).

Zheng, X. 2013. A Tutorial for the R Package SNPRelate. University of Washington, Washington, USA

Zuellig, M. P., & Sweigart, A. L. 2018. A two-locus hybrid incompatibility is widespread, polymorphic, and active in natural populations of *Mimulus*. Evolution, 72(11), 2394–2405.

Zuellig, M. P., & Sweigart, A. L. 2018. Gene duplicates cause hybrid lethality between sympatric species of *Mimulus*. PLoS genetics, 14(4)

