## Supplementary Information for "Evolution of multiple postzygotic barriers between species of the *Mimulus tilingii* complex"

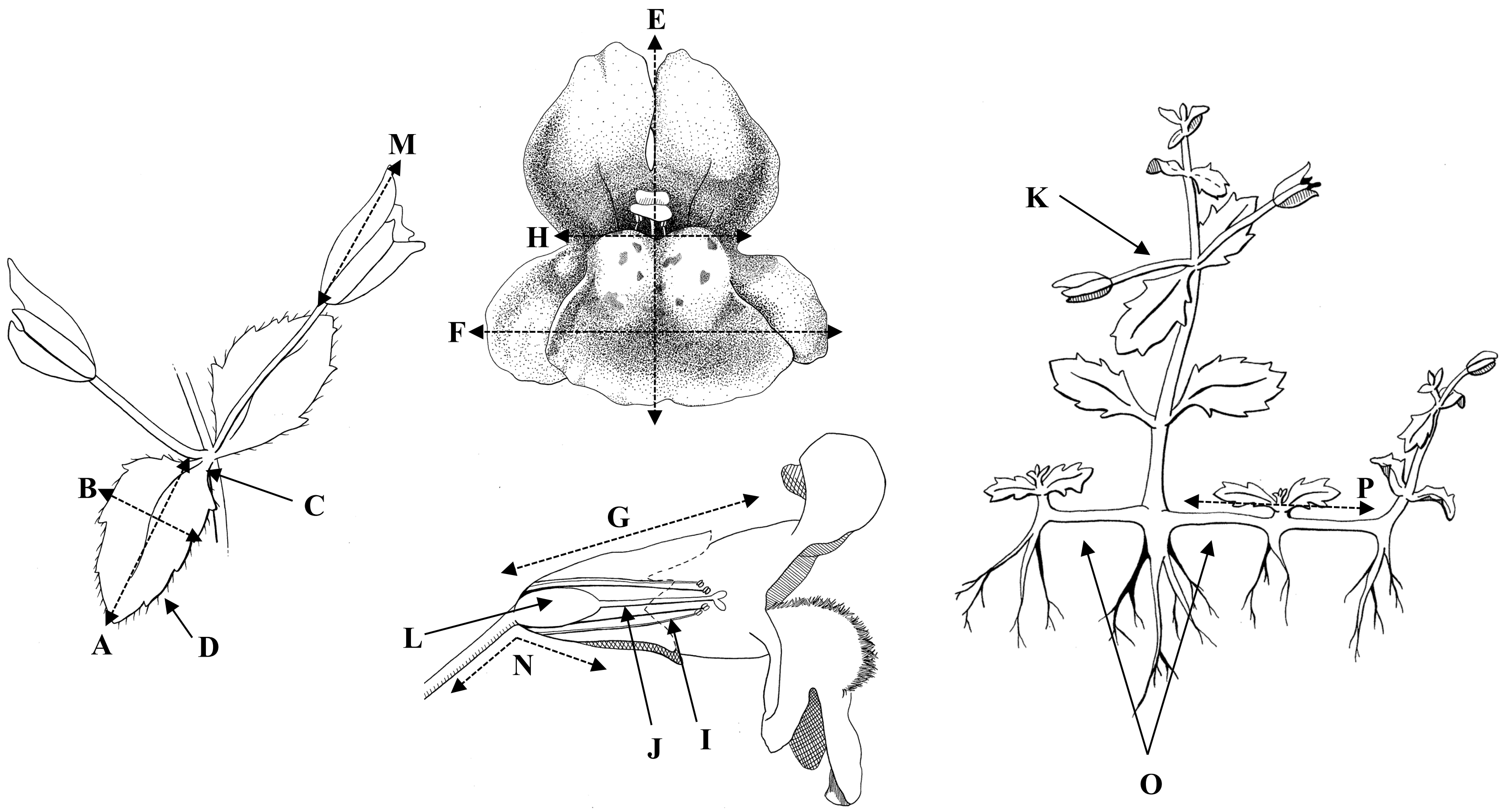


**Figure S1**. Morphological traits measured in a common garden experiment. Vegetative leaf traits measured on second leaf pair: **A.** Leaf length. **B.** Leaf width. **C.** Petiole length. **D.** Trichomes exerted past edge of leaf. **E.** Corolla height. **F.** Corolla width. **G.** Corolla tube length. **H.** Corolla tube width. **I.** Stamen length **J.** Pistil length **K.** Pedicel length **L.** Capsule length **M.** Calyx length **N.** Degree of flower nodding **O.** Number of stolons **P.** Stolon length.

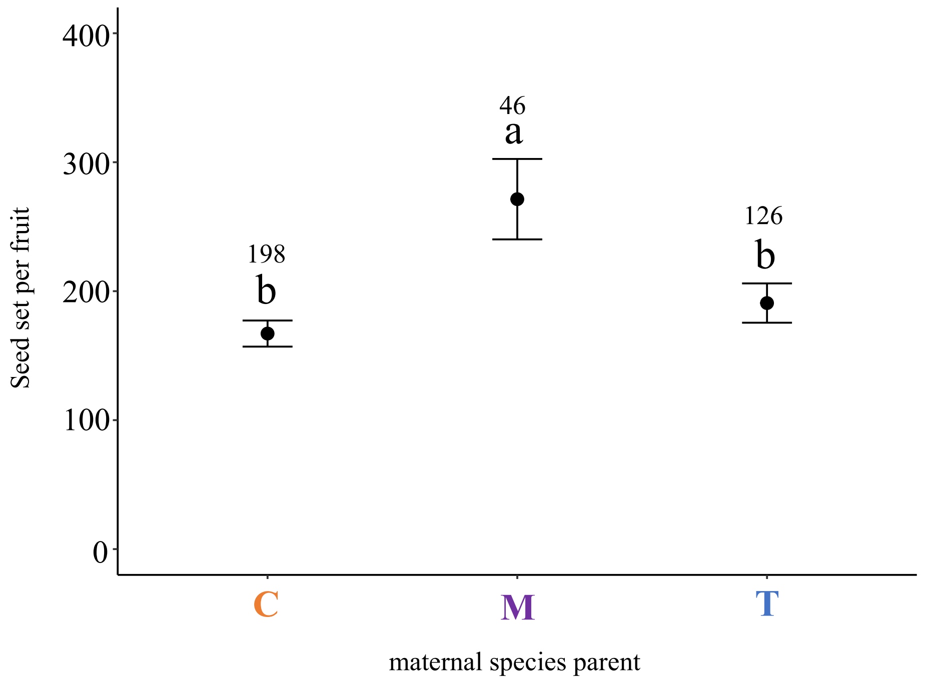


Figure S2. Total seed produced by maternal species parent of intra- and interspecific crosses when *M. caespitosa* (C), *M. minor* (M), and *M. tilingii* (T) act as the maternal parent. Least square means for each cross type are given with standard error bars. Least square means denoted by a different letter indicate significant differences among cross types ($P$ <0.05) determined by post-hoc Tukey method. Sample sizes assessed for each cross type are listed above letters

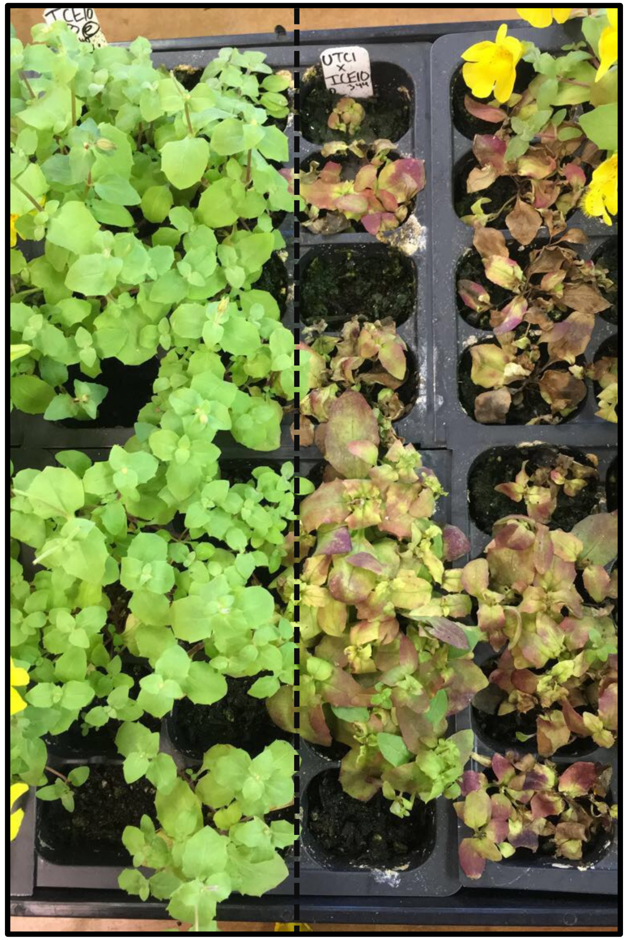


Figure S3. Hybrid inviability phenotype. Left: Intraspecific *M. tilingii* cross (TxT: ICE10xICE10). Right: Interspecific *M. caespitosa* x *M. tilingii* cross (CxT: UTC1xICE10) .

**Table S1. Description of all samples within the *Mimulus tilingii* species complex analyzed in this study.**

| Species | Population Code | Population location: Elevation (m) | State | (LAT, LONG) | Maternal Family | Generations inbred | # plants/maternal family[‡] | Crossing barriers tested? | Million Paired-End Reads | % Heterozygosity[§] | % Coverage @ 14k Sites | SRA |
|---|---|---|---|---|---|---|---|---|---|---|---|---|
| ***M. caespitosa*** | GAB | 2350 | WA | (46.88297, -121.72335) | GAB1 | 1 | 5 | X | 11.1 | 0.88 | 58.91 | |
| | | | | | GAB2 | 1 | 2 | | 14.7 | 1.09 | 86.3 | |
| ***M. caespitosa*** | KCK | 1090 | WA | (46.84036, -121.56115) | KCK1 | 1 | 5 | X | 17.2 | 1.33 | 90.41 | |
| ***M. caespitosa*** | PAG | 1880 | WA | (46.79924, -121.71255) | PAG1 | 1 | 3 | X | not sequenced | NA | NA | NA |
| | | | | | PAG2 | 1 | 5 | X | 12.8 | 1.03 | 76.92 | |
| ***M. caespitosa*** | TWN | 1594 | WA | (48.57026, -121.38164) | TWN36 | 7 | 5 | X | 15.6 | 0.53 | 85.68 | |
| ***M. caespitosa*** | UTC | 1025 | WA | (46.8004, -121.87111) | UTC1 | 1 | 5 | X | 12.7 | 0.99 | 73.86 | |
| | | | | | UTC2 | 1 | 5 | X | 12.1 | 0.75 | 75.75 | |
| ***M. minor*** | NOR | 3349 | CO | (39.015033, -107.045033) | NOR511 | 3 | 2 | X | 16.2 | 0.44 | 93.94 | |
| | | | | | NOR523 | 4 | 5 | X | 15.5 | 0.45 | 91.13 | |
| ***M. minor*** | UNP | 3728 | CO | (39.019639, -107.096053) | UNP12 | 2 | 1 | | 14.9 | 2.06 | 87.25 | |
| ***M. tilingii*** | A25 | 2343[†] | OR | (42.6364, -118.5767) | A25 | 0 | 0 | | 9.2 | 1.44 | NA | SRX6914883[¶] |
| ***M. tilingii*** | ICE | 2369 | OR | (45.13553, -117.16067) | ICE10 | 6 | 3 | X | 12.5 | 0.8 | 76.99 | |
| ***M. tilingii*** | LVR | 2751 | CA | (37.57049, -119.13544) | LVR1 | 8 | 5 | X | 54.2 | 0.66 | 99.77 | SRX1532174[¶] |
| ***M. tilingii*** | SAB | 2778 | CA | (37.12720, -118.36627) | SAB1 | 6 | 5 | X | 14 | 0.88 | 84.45 | |
| | | | | | SAB19 | 5 | 5 | | 11.6 | 0.67 | 74.23 | |
| ***M. tilingii*** | SOP | 1025 | CA | (38.266753, -119.617981) | SOP12 | 6 | 5 | X | 13.7 | 0.67 | 86.09 | |

[†]Elevation estimated from https://www.mapcoordinates.net/en

[‡]Number of individuals measured for floral and vegetative traits.

[§]Values from this column are estimated from fourfold degenerate synonymous sites

[¶]SRAs obtained from previously generated sequence data (new samples will have SRA codes entered if accepted)

**Table S2. Lsmeans and standard error (SE) for 16 morphological traits of *M. tilingii* subgroups**

| Trait | Subgroups | | | | | | Data transformation for LDA |
|---|---|---|---|---|---|---|---|
| | *M. caespitosa* | | *M. minor* | | *M. tilingii* | | |
| | Lsmeans | SE | Lsmeans | SE | Lsmeans | SE | |
| Trichomes | 24.05 | 0.94 | 1.9 | 0.49 | 344.68 | 3.34 | Normal-Quantile |
| Petiole Length (mm) | 3.48 | 0.38 | 6.74 | 1.56 | 3.08 | 0.42 | Log |
| Leaf Length (mm) | 18.05 | 0.79 | 22.91 | 2.09 | 31.02 | 1.68 | Cube-Root |
| Leaf Width (mm) | 12.16 | 0.54 | 14.14 | 1.32 | 17.57 | 0.97 | None |
| Corolla Height (mm) | 19.02 | 0.52 | 12.39 | 0.71 | 18.58 | 0.63 | None |
| Corolla Width (mm) | 16.41 | 0.45 | 14.76 | 0.85 | 21.14 | 0.72 | None |
| Tube Length (mm) | 14.6 | 0.55 | 12.95 | 1.01 | 16.65 | 0.77 | None |
| Tube Width (mm) | 10.14 | 0.29 | 10 | 0.6 | 11.74 | 0.42 | None |
| Pistil Length (mm) | 16.72 | 0.36 | 14.25 | 0.64 | 19.83 | 0.53 | None |
| Stamen Length (mm) | 13.63 | 0.31 | 13.23 | 0.62 | 15.58 | 0.43 | None |
| Pedicel Length (mm) | 17.52 | 0.91 | 9.37 | 1.01 | 21.51 | 1.37 | None |
| Capsule Length (mm) | 4.35 | 0.11 | 4.54 | 0.25 | 5.41 | 0.17 | None |
| Calyx Length (mm) | 10.68 | 0.3 | 11.98 | 0.7 | 13 | 0.45 | Log |
| Calyx Nodding (°) | 134.82 | 2 | 105.73 | 3.52 | 144.76 | 2.66 | Square-Root |
| Number of Stolons | 3.83 | 0.33 | 1.38 | 0.41 | 3.3 | 0.38 | None |
| Stolon Length (mm) | 22.93 | 1.72 | 7.05 | 1.28 | 13.85 | 1.28 | Square-Root |

Note: We fit a GLM with a gamma distribution (except trichomes and stolon branches, which used a Poisson distribution), where the trait measurements were the response variable and *M. tilingii* species was the predictor variable. We used the emmeans command in R to calculate lsmeans for the model.

**Table S3. Description of *Mimulus* maternal lines obtained from Brandvain et al. 2014)**

| Species | Population Code | Million Paired-End Reads | % heterozygosity | % Coverage @ 14k sites | SRA |
|---|---|---|---|---|---|
| ***M. dentilobus*** | DENT | 12.1 | 0.53 | 65.25 | SRX030541 |
| ***M. guttatus*** | AHQT1 | 34.9 | 0.73 | 98.83 | SRX142379 |
| ***M. guttatus*** | CACG6 | 70.2 | 0.88 | 99.62 | SRX525044 |
| ***M. guttatus*** | MED82 | 44.8 | 0.92 | 99.5 | SRA166889 |
| ***M. guttatus*** | SLP | 41.9 | 0.89 | 99.39 | SRX142377 |
| ***M. nasutus*** | CACN9 | 46.6 | 0.6 | 99.6 | SRX525048 |
| ***M. nasutus*** | NHN | 46.1 | 0.61 | 99.39 | SRX525051 |

**Table S4. Comparisons of crosses performed within maternal lines and between maternal lines of the same species within the *M. tilingii* complex.**

| Barrier | Species | Type | Lsmean | SE | Estimate | SE | Z-ratio | P-value |
|---|---|---|---|---|---|---|---|---|
| **Seed Set** | *M. caespitosa* | between-lines | 171.2 | 13.39 | -6.93E-04 | 1.28E-03 | -0.54 | 0.59 |
| | | within-lines | 153.06 | 28.08 | | | | |
| | *M. minor* | between-lines | 337.5 | 119.11 | -1.63E-04 | 1.26E-03 | -0.13 | 0.9 |
| | | within-lines | 319.8 | 71.38 | | | | |
| | *M. tilingii* | between-lines | 165.22 | 18.78 | 9.70E-04 | 9.92E-04 | 0.98 | 0.33 |
| | | within-lines | 196.73 | 27.7 | | | | |
| **Visual Seed Assessment** | *M. caespitosa* | between-lines | 0.94 | 0.01 | 1.77 | 0.13 | 7.66 | <0.001 |
| | | within-lines | 0.9 | 0.02 | | | | |
| | *M. minor* | between-lines | 0.93 | 0.03 | 0.96 | 0.2 | -0.22 | 0.82 |
| | | within-lines | 0.93 | 0.03 | | | | |
| | *M. tilingii* | between-lines | 0.91 | 0.02 | 1.5 | 0.13 | 4.8 | <0.001 |
| | | within-lines | 0.87 | 0.02 | | | | |
| **Germination Success** | *M. caespitosa* | between-lines | 0.56 | 0.07 | 1.47 | 0.1 | 6 | <0.001 |
| | | within-lines | 0.46 | 0.07 | | | | |
| | *M. minor* | between-lines | 0.68 | 0.06 | 19.36 | 5.85 | 9.8 | <0.001 |
| | | within-lines | 0.1 | 0.02 | | | | |
| | *M. tilingii* | between-lines | 0.63 | 0.13 | 0.94 | 0.14 | -0.39 | 0.7 |
| | | within-lines | 0.64 | 0.13 | | | | |
| **F1 Seed Set** | *M. caespitosa* | between-lines | 214.96 | 1.89 | 6.48E-05 | 7.96E-04 | 0.081 | 0.94 |
| | | within-lines | 218 | 3.58 | | | | |
| | *M. minor* | between-lines | 511.5 | 67.37 | -1.40E-03 | 5.57E-04 | -2.512 | 0.01 |
| | | within-lines | 298.13 | 43.9 | | | | |
| | *M. tilingii* | between-lines | 234.28 | 2.95 | -8.71E-04 | 1.34E-03 | -0.648 | 0.52 |
| | | within-lines | 194.57 | 5.27 | | | | |
| **F1 Pollen Viability** | *M. caespitosa* | between-lines | 0.96 | 0.02 | 4.06 | 0.49 | 11.55 | <0.001 |
| | | within-lines | 0.87 | 0.07 | | | | |
| | *M. minor* | between-lines | 0.95 | 0.01 | 1.38 | 0.23 | 1.94 | 0.053 |
| | | within-lines | 0.93 | 0.01 | | | | |
| | *M. tilingii* | between-lines | 0.73 | 0.06 | 0.38 | 0.02 | -17.54 | <0.001 |
| | | within-lines | 0.88 | 0.03 | | | | |

**Table S5. Crosses performed within and between populations and species of the *Mimulus tilingii* complex**

| Unique Crosses† | Number of fruits per cross‡ | Number of fruits assayed for seed germination§ |
|---|---|---|
| **CxC** | **116** | **54** |
| GAB1xGAB1 | 2 | 2 |
| GAB1xKCK1 | 3 | 1 |
| GAB1xPAG1 | 2 | |
| GAB1xTWN36 | 2 | |
| GAB1xUTC1 | 3 | 1 |
| GAB1xUTC2 | 2 | 2 |
| KCK1xGAB1 | 1 | 1 |
| KCK1xKCK1 | 4 | 2 |
| KCK1xPAG1 | 1 | |
| KCK1xPAG2 | 3 | 2 |
| KCK1xTWN36 | 5 | 2 |
| KCK1xUTC1 | 2 | 1 |
| KCK1xUTC2 | 1 | 1 |
| PAG1xKCK1 | 1 | 1 |
| PAG1xPAG1 | 1 | |
| PAG1xTWN36 | 1 | |
| PAG1xUTC2 | 1 | 1 |
| PAG2xKCK1 | 4 | 1 |
| PAG2xPAG1 | 3 | 1 |
| PAG2xPAG2 | 2 | 2 |
| PAG2xTWN36 | 5 | 1 |
| PAG2xUTC1 | 6 | 2 |
| PAG2xUTC2 | 2 | |
| TWN36xGAB1 | 2 | 1 |
| TWN36xKCK1 | 3 | 1 |
| TWN36xPAG1 | 2 | 1 |
| TWN36xPAG2 | 3 | 1 |
| TWN36xTWN36 | 5 | 2 |
| TWN36xUTC1 | 4 | 2 |
| TWN36xUTC2 | 4 | 1 |
| UTC1xGAB1 | 2 | 1 |
| UTC1xKCK1 | 4 | 1 |
| UTC1xPAG1 | 1 | |
| UTC1xPAG2 | 1 | 1 |
| UTC1xTWN36 | 2 | 2 |
| UTC1xUTC1 | 4 | 2 |
| UTC1xUTC2 | 2 | 2 |
| UTC2xGAB1 | 3 | 3 |
| UTC2xKCK1 | 3 | 1 |
| UTC2xPAG1 | 1 | |
| UTC2xPAG2 | 4 | 2 |
| UTC2xTWN36 | 3 | 2 |
| UTC2xUTC1 | 3 | 2 |
| UTC2xUTC2 | 3 | 2 |
| **CxM** | **22** | **13** |
| GAB1xNOR511 | 1 | 1 |
| KCK1xNOR511 | 1 | |
| PAG1xNOR511 | 2 | 1 |

| | | |
|---|---|---|
| PAG2xNOR511 | 3 | 1 |
| PAG2xNOR523 | 1 | 1 |
| TWN36xNOR511 | 3 | 2 |
| TWN36xNOR523 | 3 | 2 |
| UTC1xNOR511 | 2 | |
| UTC2xNOR511 | 4 | 3 |
| UTC2xNOR523 | 2 | 2 |
| **MxC** | **25** | **13** |
| NOR511xGAB1 | 1 | |
| NOR511xKCK1 | 3 | 3 |
| NOR511xPAG1 | 2 | 1 |
| NOR511xPAG2 | 2 | |
| NOR511xTWN36 | 1 | |
| NOR511xUTC1 | 2 | 1 |
| NOR511xUTC2 | 4 | |
| NOR523xGAB1 | 2 | 2 |
| NOR523xKCK1 | 2 | 2 |
| NOR523xTWN36 | 2 | 2 |
| NOR523xUTC1 | 2 | 2 |
| NOR523xUTC2 | 2 | |
| **MxM** | **7** | **3** |
| NOR511xNOR511 | 2 | 1 |
| NOR511xNOR523 | 1 | |
| NOR523xNOR511 | 1 | 1 |
| NOR523xNOR523 | 3 | 1 |
| **MxT** | **14** | **9** |
| NOR511xICE10 | 3 | 1 |
| NOR511xLVR | 1 | 1 |
| NOR511xSAB | 1 | |
| NOR511xSOP12 | 1 | 1 |
| NOR523xICE10 | 2 | 1 |
| NOR523xLVR | 2 | 2 |
| NOR523xSAB | 2 | 2 |
| NOR523xSOP12 | 2 | 1 |
| **TxM** | **14** | **11** |
| ICE10xNOR511 | 3 | 3 |
| ICE10xNOR523 | 1 | 1 |
| LVRxNOR511 | 3 | 1 |
| LVRxNOR523 | 2 | 2 |
| SABxNOR511 | 2 | 2 |
| SOP12xNOR511 | 2 | 1 |
| SOP12xNOR523 | 1 | 1 |
| **TxT** | **37** | **19** |
| ICE10xICE10 | 4 | 2 |
| ICE10xLVR | 3 | 2 |
| ICE10xSOP12 | 3 | 3 |
| LVRxLVR | 5 | 3 |
| LVRxSAB | 3 | |
| LVRxSOP12 | 1 | |
| SABxICE10 | 4 | 3 |
| SABxLVR | 2 | |
| SABxSAB | 2 | 2 |
| SABxSOP12 | 2 | |
| SOP12xICE10 | 2 | 1 |
| SOP12xLVR | 2 | 1 |
| SOP12xSAB | 1 | |

| | | |
|---|---|---|
| SOP12xSOP12 | 3 | 1 |
| **TxC** | **75** | **40** |
| ICE10xGAB1 | 1 | 1 |
| ICE10xKCK1 | 3 | 3 |
| ICE10xPAG1 | 2 | 2 |
| ICE10xPAG2 | 2 | 2 |
| ICE10xTWN36 | 3 | 1 |
| ICE10xUTC1 | 3 | 2 |
| ICE10xUTC2 | 2 | 2 |
| LVRxGAB1 | 1 | |
| LVRxKCK1 | 1 | |
| LVRxTWN36 | 7 | 2 |
| LVRxUTC1 | 1 | 1 |
| LVRxUTC2 | 2 | 4 |
| SABxGAB1 | 2 | 1 |
| SABxKCK1 | 3 | 1 |
| SABxPAG2 | 8 | 3 |
| SABxTWN36 | 2 | |
| SABxUTC1 | 4 | 4 |
| SABxUTC2 | 2 | 1 |
| SOP12xGAB1 | 3 | 2 |
| SOP12xKCK1 | 5 | 1 |
| SOP12xPAG1 | 5 | 4 |
| SOP12xPAG2 | 1 | |
| SOP12xTWN36 | 5 | |
| SOP12xUTC1 | 5 | 2 |
| SOP12xUTC2 | 2 | 1 |
| **CxT** | **60** | **47** |
| GAB1xICE10 | 2 | 1 |
| GAB1xLVR | 1 | 1 |
| GAB1xSAB | 2 | 1 |
| GAB1xSOP12 | 1 | 1 |
| KCK1xICE10 | 3 | 3 |
| KCK1xLVR | 2 | 2 |
| KCK1xSAB | 3 | 3 |
| KCK1xSOP12 | 2 | 2 |
| PAG1xICE10 | 1 | 1 |
| PAG2xICE10 | 2 | 1 |
| PAG2xLVR | 1 | 1 |
| PAG2xSOP12 | 4 | 4 |
| TWN36xICE10 | 3 | 1 |
| TWN36xLVR | 4 | 2 |
| TWN36xSAB | 2 | |
| TWN36xSOP12 | 3 | 1 |
| UTC1xICE10 | 3 | 3 |
| UTC1xLVR | 2 | 2 |
| UTC1xSAB | 2 | 1 |
| UTC1xSOP12 | 3 | 3 |
| UTC2xICE10 | 4 | 4 |
| UTC2xLVR | 4 | 4 |
| UTC2xSAB | 3 | 2 |
| UTC2xSOP12 | 3 | 3 |

†Unique cross combinations of maternal families, female parent listed first.

‡Numbers of fruits listed in this column were used to assess postmating,

prezygotic isolation and hybrid seed inviability.

[§]Numbers of fruits listed in this column were used to assess seed germination rates.

**Table S6. Assessment of hybrid inviability and hybrid sterility between species of the *Mimulus tilingii* complex.**

| Unique Crosses | No. F1 seedlings tansplanted per cross | No. F1s that flowered | Proportion flowering progeny | Proportion necrotic progeny | Average days to flower (from seedling) | Individuals with pollen viability measured† | No. F1 individuals used for backcross | Pollen donor | Total F1 crosses |
|---|---|---|---|---|---|---|---|---|---|
| **CxC** | **199** | **199** | **1** | **0** | **49.56** | **136** | **76** | | **95** |
| GAB1xGAB1 | 15 | 15 | 1 | 0 | 50.13 | 10 | 2 | GAB1 | 3 |
| KCK1xKCK1 | 10 | 10 | 1 | 0 | 49.7 | 7 | 3 | KCK1 | 3 |
| KCK1xPAG2 | 16 | 16 | 1 | 0 | 42.75 | 11 | 9 | PAG2 | 10 |
| KCK1xUTC1 | 8 | 8 | 1 | 0 | 57 | 4 | 3 | KCK1 | 4 |
| PAG1xTWN36 | 10 | 10 | 1 | 0 | 51.4 | 7 | 4 | TWN36 | 5 |
| PAG2xPAG2 | 4 | 4 | 1 | 0 | 57.5 | 2 | 1 | PAG2 | 2 |
| PAG2xUTC1 | 15 | 15 | 1 | 0 | 54.53 | 7 | 7 | UTC1 | 8 |
| PAG1xUTC2 | 16 | 16 | 1 | 0 | 53.31 | 8 | 6 | UTC2 | 8 |
| UTC1xPAG2 | 10 | 10 | 1 | 0 | 42.5 | 8 | 4 | UTC1 | 5 |
| UTC1xUTC1 | 12 | 12 | 1 | 0 | 56.75 | 10 | 3 | UTC1 | 4 |
| UTC2xPAG2 | 11 | 11 | 1 | 0 | 38.27 | 8 | 5 | PAG2 | 6 |
| UTC2xUTC1 | 6 | 6 | 1 | 0 | 63.67 | 6 | 3 | UTC2 | 5 |
| UTC2xUTC2 | 11 | 11 | 1 | 0 | 49.45 | 8 | 5 | UTC2 | 5 |
| TWN36xPAG1 | 16 | 16 | 1 | 0 | 46.13 | 14 | 6 | TWN36 | 8 |
| TWN36xPAG2 | 12 | 12 | 1 | 0 | 50.8 | 11 | 5 | PAG2 | 7 |
| TWN36xTWN36 | 15 | 15 | 1 | 0 | 46.73 | 7 | 3 | TWN36 | 4 |
| TWN36xUTC1 | 12 | 12 | 1 | 0 | 31.9 | 17 | 7 | UTC1 | 8 |
| **CxM** | **57** | **57** | **1** | **0** | **36.23** | **38** | **22** | | **27** |
| PAG1xNOR511 | 12 | 12 | 1 | 0 | 35.5 | 8 | 4 | NOR511 | 5 |
| PAG2xNOR511 | 15 | 15 | 1 | 0 | 36.13 | 8 (2) | 5 | PAG2 | 6 |
| TWN36xNOR511 | 7 | 7 | 1 | 0 | 36.86 | 4 | 4 | TWN36 | 5 |
| UTC2xNOR511 | 13 | 13 | 1 | 0 | 35.54 | 9 | 4 | UTC2 | 5 |
| UTC2xNOR523 | 10 | 10 | 1 | 0 | 37.1 | 9 | 5 | UTC2 | 6 |
| **MxC** | **56** | **56** | **1** | **0** | **47.64** | **34** | **17** | | **26** |
| NOR511xKCK1 | 16 | 16 | 1 | 0 | 46.63 | 13 | 6 | NOR511/KCK1 | 11 |
| NOR511xPAG1 | 16 | 16 | 1 | 0 | 47.81 | 10 | 3 | NOR511/PAG1 | 5 |
| NOR511xUTC1 | 9 | 9 | 1 | 0 | 58.33 | 4 | 4 | UTC1 | 6 |
| NOR523xGAB1 | 15 | 15 | 1 | 0 | 37.8 | 7 (1) | 4 | GAB1 | 4 |
| **MxM** | **23** | **23** | **1** | **0** | **42.07** | **13** | **9** | | **12** |
| NOR511xNOR511 | 11 | 11 | 1 | 0 | 47.64 | 7 | 4 | NOR511 | 6 |
| NOR523xNOR511 | 12 | 12 | 1 | 0 | 36.5 | 6 | 5 | NOR511 | 6 |
| **TxM** | **54** | **54** | **1** | **0** | **43.4** | **26** | **11** | | **12** |
| ICE10xNOR511 | 13 | 13 | 1 | 0 | 45.31 | 7 (4) | 3 | ICE10 | 3 |
| SAB1xNOR511 | 23 | 23 | 1 | 0 | 44.7 | 9 | 3 | NOR511 | 4 |
| SAB1xNOR523 | 3 | 3 | 1 | 0 | 42 | 3 | 3 | SAB1 | 3 |
| SOP12xNOR511 | 15 | 15 | 1 | 0 | 41.6 | 7 | 2 | NOR511 | 2 |
| **TxT** | **93** | **93** | **1** | **0** | **45.06** | **57** | **34** | | **38** |
| ICE10xICE10 | 11 | 11 | 1 | 0 | 61.91 | 7 | 2 | ICE10 | 2 |
| ICE10xLVR1 | 10 | 10 | 1 | 0 | 36.2 | 8 | 7 | LVR1 | 7 |
| ICE10xSOP12 | 15 | 15 | 1 | 0 | 42.47 | 7 | 3 | SOP12 | 4 |
| LVR1xLVR1 | 16 | 16 | 1 | 0 | 51.5 | 7 | 2 | LVR1 | 2 |
| LVR1xSOP12 | 5 | 5 | 1 | 0 | 36 | 5 | 4 | SOP12 | 5 |
| SOP12xICE10 | 12 | 12 | 1 | 0 | 32.25 | 8 | 7 | SOP12 | 9 |
| SOP12xLVR1 | 11 | 11 | 1 | 0 | 45.18 | 6 | 6 | SOP12 | 6 |
| SOP12xSOP12 | 13 | 13 | 1 | 0 | 55 | 9 | 3 | SOP12 | 3 |
| **TxC** | **107** | **93** | **0.87** | **0.13** | **43.44** | **65** | **43** | | **63** |
| ICE10xGAB1 | 16 | 8 | 0.5 | 0.5 | 49.13 | 7 | 2 | ICE10 | 3 |
| ICE10xKCK1 | 16 | 16 | 1 | 0 | 43 | 8 | 5 | KCK1 | 6 |
| ICE10xPAG2 | 13 | 13 | 1 | 0 | 49 | 7 | 4 | PAG2 | 6 |
| ICE10xTWN36 | 13 | 13 | 1 | 0 | 48.64 | 8 | 7 | TWN36 | 10 |
| ICE10xUTC1 | 11 | 5 | 0.45 | 0.55 | 52.2 | 4 | 5 | UTC1 | 12 |
| LVR1xUTC2 | 12 | 12 | 1 | 0 | 31.58 | 8 | 6 | LVR1 | 6 |
| SAB1xPAG2 | 8 | 8 | 1 | 0 | 38.88 | 8 | 4 | PAG2 | 7 |
| SAB1xUTC1 | 8 | 8 | 1 | 0 | 37.13 | 8 | 5 | UTC1 | 6 |
| SAB1xUTC2 | 10 | 10 | 1 | 0 | 41.4 | 7 | 5 | UTC2 | 7 |
| **CxT** | **60** | **44** | **0.73** | **0.27** | **51.38** | **29** | **19** | | **24** |
| GAB1xICE10 | 8 | 6 | 0.75 | 0.25 | 44.67 | 6 | 3 | GAB1 | 4 |
| GAB1xLVR1 | 16 | 16 | 1 | 0 | 45.44 | 7 (2) | 7 | LVR1 | 8 |
| KCK1xSAB1 | 11 | 11 | 1 | 0 | 67.09 | 5 (5) | 4 | KCK1 | 5 |
| PAG2xSOP12 | 3 | 3 | 1 | 0 | 52.67 | 3 | 1 | KCK1 | 2 |
| TWN36xLVR1 | 1 | 1 | 1 | 0 | 56 | 1 | -- | -- | -- |
| UTC1xICE10 | 14 | 0 | 0 | 1 | -- | -- | -- | -- | -- |
| UTC2xLVR1 | 7 | 7 | 1 | 0 | 42.43 | 7 | 4 | LVR1 | 5 |
| **Total** | **649** | **619** | **--** | **--** | **--** | **399** | **231** | **--** | **297** |

†In parentheses are the number of individuals with anthers that were either shriveled and/or produced no pollen

**Table S7. Trait coefficients for each linear discriminant. Largest coefficient (positive or negative) indicate traits that contribute most to that discriminant function.**

| Trait | LD1 | LD2 | Total † |
|---|---|---|---|
| Trichomes | 0.99 | 0.34 | 1.32 |
| Petiole Length | -0.23 | -0.35 | 0.58 |
| Leaf Length | 1.32 | -0.59 | 1.91 |
| Leaf Width | -0.17 | 0.56 | 0.73 |
| Corolla Height | -0.83 | 0.49 | 1.32 |
| Corolla Width | 0.35 | -0.43 | 0.78 |
| Tube Length | -0.44 | 0.22 | 0.66 |
| Tube Width | -0.11 | -0.31 | 0.42 |
| Pistil Length | 0.01 | 1.63 | 1.64 |
| Stamen Length | -0.03 | -0.56 | 0.6 |
| Pedicel Length | -0.47 | 0.34 | 0.81 |
| Capsule Length | 0.5 | -0.74 | 1.24 |
| Calyx Length | 0.62 | 0.1 | 0.72 |
| Calyx Nodding | 0.65 | 1 | 1.65 |
| Stolon Branches | 0.13 | 0.49 | 0.63 |
| Stolon Branch Length | -0.62 | 0.16 | 0.78 |

†Absolute value of LD1 and LD2

**Table S8. LDA model prediction based on probability that individual belongs to a given class**

| Maternal line | Individual | Proposed Species | Class Probability | | |
|---|---|---|---|---|---|
| | | | *M. caespitosa* | *M. minor* | *M. tilingii* |
| **GAB1** | 1 | C | 9.77E-01 | 7.76E-09 | 2.28E-02 |
| **GAB1** | 2 | C | 1.00E+00 | 2.21E-15 | 1.09E-07 |
| **GAB1** | 3 | C | 1.00E+00 | 7.29E-09 | 3.00E-06 |
| **GAB1** | 4 | C | 9.95E-01 | 2.40E-12 | 4.63E-03 |
| **GAB1** | 5 | C | 1.00E+00 | 2.15E-07 | 4.86E-05 |
| **GAB2** | 1 | C | 1.00E+00 | 1.45E-15 | 8.97E-09 |
| **GAB2** | 2 | C | 1.00E+00 | 3.31E-12 | 1.44E-07 |
| **KCK1** | 1 | C | 1.00E+00 | 2.56E-10 | 5.69E-08 |
| **KCK1** | 2 | C | 1.00E+00 | 4.50E-09 | 4.51E-06 |
| **KCK1** | 3 | C | 1.00E+00 | 1.60E-13 | 2.63E-08 |
| **KCK1** | 4 | C | 1.00E+00 | 1.80E-13 | 1.84E-09 |
| **KCK1** | 5 | C | 1.00E+00 | 8.49E-14 | 1.51E-06 |
| **PAG1** | 1 | C | 1.00E+00 | 1.14E-10 | 3.39E-08 |
| **PAG1** | 2 | C | 1.00E+00 | 9.98E-16 | 1.40E-06 |
| **PAG1** | 3 | C | 1.00E+00 | 6.35E-13 | 1.86E-11 |
| **PAG2** | 1 | C | 9.93E-01 | 5.27E-11 | 7.42E-03 |
| **PAG2** | 2 | C | 1.00E+00 | 3.49E-12 | 8.50E-10 |
| **PAG2** | 3 | C | 1.00E+00 | 1.12E-14 | 2.51E-07 |
| **PAG2** | 4 | C | 1.00E+00 | 8.67E-09 | 5.64E-09 |
| **PAG2** | 5 | C | 1.00E+00 | 3.84E-13 | 9.75E-08 |
| **TWN36** | 1 | C | 1.00E+00 | 2.14E-12 | 4.62E-06 |
| **TWN36** | 2 | C | 1.00E+00 | 2.68E-06 | 3.52E-04 |
| **TWN36** | 3 | C | 1.00E+00 | 3.12E-04 | 1.10E-05 |
| **TWN36** | 4 | C | 9.99E-01 | 2.68E-06 | 7.78E-04 |
| **TWN36** | 5 | C | 9.99E-01 | 2.65E-09 | 5.09E-04 |
| **UTC1** | 1 | C | 1.00E+00 | 5.84E-10 | 1.47E-11 |
| **UTC1** | 2 | C | 1.00E+00 | 9.74E-14 | 5.04E-14 |
| **UTC1** | 3 | C | 1.00E+00 | 6.33E-08 | 2.03E-09 |
| **UTC1** | 4 | C | 1.00E+00 | 6.84E-09 | 4.46E-09 |
| **UTC1** | 5 | C | 1.00E+00 | 7.64E-13 | 1.98E-08 |
| **UTC2** | 1 | C | 1.00E+00 | 1.41E-09 | 4.27E-08 |
| **UTC2** | 2 | C | 1.00E+00 | 7.15E-11 | 1.30E-08 |
| **UTC2** | 3 | C | 1.00E+00 | 1.10E-15 | 3.05E-08 |
| **UTC2** | 4 | C | 1.00E+00 | 3.45E-11 | 2.62E-10 |
| **UTC2** | 5 | C | 1.00E+00 | 1.11E-09 | 4.20E-07 |
| **NOR511** | 1 | M | 1.84E-10 | 1.00E+00 | 1.44E-14 |
| **NOR511** | 2 | M | 3.11E-11 | 1.00E+00 | 3.46E-13 |
| **NOR523** | 1 | M | 7.21E-12 | 1.00E+00 | 7.24E-14 |
| **NOR523** | 2 | M | 2.56E-11 | 1.00E+00 | 7.40E-11 |

| | | | | | |
|---|---|---|---|---|---|
| **NOR523** | 3 | M | 2.80E-05 | 1.00E+00 | 9.73E-06 |
| **NOR523** | 4 | M | 1.87E-08 | 1.00E+00 | 1.38E-05 |
| **NOR523** | 5 | M | 3.93E-11 | 1.00E+00 | 5.66E-11 |
| **UNP12** | 3 | M | 7.40E-09 | 1.00E+00 | 3.71E-11 |
| **ICE10** | 1 | T | 1.77E-06 | 4.22E-13 | 1.00E+00 |
| **ICE10** | 2 | T | 1.61E-04 | 2.09E-06 | 1.00E+00 |
| **ICE10** | 3 | T | 8.35E-04 | 7.76E-13 | 9.99E-01 |
| **LVR1** | 1 | T | 1.98E-06 | 2.67E-13 | 1.00E+00 |
| **LVR1** | 2 | T | 3.31E-02 | 1.99E-11 | 9.67E-01 |
| **LVR1** | 3 | T | 3.93E-08 | 1.76E-14 | 1.00E+00 |
| **LVR1** | 4 | T | 1.49E-10 | 5.86E-15 | 1.00E+00 |
| **LVR1** | 5 | T | 7.28E-07 | 4.95E-13 | 1.00E+00 |
| **SAB1** | 1 | T | 8.12E-11 | 2.52E-13 | 1.00E+00 |
| **SAB1** | 2 | T | 1.30E-08 | 2.75E-11 | 1.00E+00 |
| **SAB1** | 3 | T | 2.12E-08 | 7.46E-14 | 1.00E+00 |
| **SAB1** | 4 | T | 3.03E-07 | 5.31E-13 | 1.00E+00 |
| **SAB1** | 5 | T | 2.86E-08 | 2.38E-09 | 1.00E+00 |
| **SAB19** | 1 | T | 4.72E-06 | 3.34E-18 | 1.00E+00 |
| **SAB19** | 2 | T | 1.06E-09 | 1.04E-10 | 1.00E+00 |
| **SAB19** | 3 | T | 3.67E-06 | 7.76E-11 | 1.00E+00 |
| **SAB19** | 4 | T | 2.92E-07 | 2.58E-09 | 1.00E+00 |
| **SAB19** | 5 | T | 1.06E-04 | 2.38E-12 | 1.00E+00 |
| **SOP12** | 1 | T | 3.90E-08 | 4.97E-09 | 1.00E+00 |
| **SOP12** | 2 | T | 8.91E-08 | 4.50E-09 | 1.00E+00 |
| **SOP12** | 3 | T | 1.05E-06 | 1.30E-10 | 1.00E+00 |
| **SOP12** | 4 | T | 5.25E-09 | 5.37E-11 | 1.00E+00 |
| **SOP12** | 5 | T | 4.34E-05 | 1.08E-06 | 1.00E+00 |

**Table S9. Pairwise nucleotide diversity comparisons among *Mimulus* species (C = *M. caespitosa* , M = *M. minor* , T = *M. tilingii* , G = *M. guttatus* , N= *M. nasutus* )**

| Sample 1 | Sample 2 | Comparison | Intra-population? | No. sites | No. polymorphic sites | Dxy/pi |
|---|---|---|---|---|---|---|
| GAB1 | GAB2 | C x C | X | 478593 | 5711 | 0.012 |
| GAB1 | KCK1 | C x C | | 542249 | 6291 | 0.012 |
| GAB1 | PAG2 | C x C | | 379433 | 5205 | 0.014 |
| GAB1 | TWN36 | C x C | | 479699 | 6428 | 0.013 |
| GAB1 | UTC1 | C x C | | 346059 | 4742 | 0.014 |
| GAB1 | UTC2 | C x C | | 363270 | 4920 | 0.014 |
| GAB2 | KCK1 | C x C | | 1179245 | 11391 | 0.01 |
| GAB2 | PAG2 | C x C | | 792455 | 8692 | 0.011 |
| GAB2 | TWN36 | C x C | | 1044661 | 12428 | 0.012 |
| GAB2 | UTC1 | C x C | | 719855 | 8238 | 0.011 |
| GAB2 | UTC2 | C x C | | 764229 | 8251 | 0.011 |
| KCK1 | PAG2 | C x C | | 926215 | 9325 | 0.01 |
| KCK1 | TWN36 | C x C | | 1208062 | 13666 | 0.011 |
| KCK1 | UTC1 | C x C | | 843450 | 7498 | 0.009 |
| KCK1 | UTC2 | C x C | | 885378 | 8327 | 0.009 |
| PAG2 | TWN36 | C x C | | 815627 | 10539 | 0.013 |
| PAG2 | UTC1 | C x C | | 579476 | 6653 | 0.011 |
| PAG2 | UTC2 | C x C | | 608203 | 6698 | 0.011 |
| TWN36 | UTC1 | C x C | | 739034 | 9475 | 0.013 |
| TWN36 | UTC2 | C x C | | 780572 | 9606 | 0.012 |
| UTC1 | UTC2 | C x C | X | 552896 | 5794 | 0.01 |
| NOR511 | NOR523 | M x M | X | 1892185 | 4682 | 0.002 |
| NOR511 | UNP12 | M x M | | 1617266 | 16895 | 0.01 |
| NOR523 | UNP12 | M x M | | 1467315 | 15726 | 0.011 |
| A25 | ICE10 | T x T | | 86786 | 2927 | 0.034 |
| A25 | LVR | T x T | | 152911 | 5679 | 0.037 |
| A25 | SAB | T x T | | 101043 | 3691 | 0.037 |
| A25 | SAB19 | T x T | | 80550 | 3016 | 0.037 |
| A25 | SOP12 | T x T | | 104890 | 3616 | 0.034 |
| ICE10 | LVR | T x T | | 1466657 | 43164 | 0.029 |
| ICE10 | SAB | T x T | | 848641 | 28434 | 0.034 |
| ICE10 | SAB19 | T x T | | 654113 | 22176 | 0.034 |
| ICE10 | SOP12 | T x T | | 889458 | 23633 | 0.027 |
| LVR | SAB | T x T | | 1836031 | 54519 | 0.03 |
| LVR | SAB19 | T x T | | 1416846 | 42896 | 0.03 |
| LVR | SOP12 | T x T | | 1918602 | 50227 | 0.026 |

| | | | | | | |
|---|---|---|---|---|---|---|
| SAB | SAB19 | T x T | X | 832066 | 12603 | 0.015 |
| SAB | SOP12 | T x T | | 1088685 | 34531 | 0.032 |
| SAB19 | SOP12 | T x T | | 841431 | 27514 | 0.033 |
| GAB1 | NOR511 | C x M | | 607861 | 25591 | 0.042 |
| GAB1 | NOR523 | C x M | | 551162 | 23136 | 0.042 |
| GAB1 | UNP12 | C x M | | 486262 | 16734 | 0.034 |
| GAB2 | NOR511 | C x M | | 1385590 | 55454 | 0.04 |
| GAB2 | NOR523 | C x M | | 1253335 | 50202 | 0.04 |
| GAB2 | UNP12 | C x M | | 1093456 | 35142 | 0.032 |
| KCK1 | NOR511 | C x M | | 1622882 | 66563 | 0.041 |
| KCK1 | NOR523 | C x M | | 1464066 | 60192 | 0.041 |
| KCK1 | UNP12 | C x M | | 1271905 | 42747 | 0.034 |
| PAG2 | NOR511 | C x M | | 1076027 | 44631 | 0.041 |
| PAG2 | NOR523 | C x M | | 972227 | 40381 | 0.042 |
| PAG2 | UNP12 | C x M | | 847868 | 28851 | 0.034 |
| TWN36 | NOR511 | C x M | | 1434096 | 59212 | 0.041 |
| TWN36 | NOR523 | C x M | | 1296349 | 53658 | 0.041 |
| TWN36 | UNP12 | C x M | | 1120902 | 38441 | 0.034 |
| UTC1 | NOR511 | C x M | | 965764 | 40341 | 0.042 |
| UTC1 | NOR523 | C x M | | 877295 | 36740 | 0.042 |
| UTC1 | UNP12 | C x M | | 764191 | 26224 | 0.034 |
| UTC2 | NOR511 | C x M | | 1022333 | 42498 | 0.042 |
| UTC2 | NOR523 | C x M | | 925376 | 38654 | 0.042 |
| UTC2 | UNP12 | C x M | | 807475 | 27551 | 0.034 |
| GAB1 | A25 | C x T | | 52916 | 2229 | 0.042 |
| GAB1 | ICE10 | C x T | | 372174 | 16491 | 0.044 |
| GAB1 | LVR | C x T | | 758041 | 34179 | 0.045 |
| GAB1 | SAB | C x T | | 454908 | 19571 | 0.043 |
| GAB1 | SAB19 | C x T | | 350240 | 15664 | 0.045 |
| GAB1 | SOP12 | C x T | | 471532 | 20998 | 0.045 |
| GAB2 | A25 | C x T | | 95464 | 4050 | 0.042 |
| GAB2 | ICE10 | C x T | | 811522 | 34532 | 0.043 |
| GAB2 | LVR | C x T | | 1757250 | 76026 | 0.043 |
| GAB2 | SAB | C x T | | 1000513 | 41712 | 0.042 |
| GAB2 | SAB19 | C x T | | 766543 | 33065 | 0.043 |
| GAB2 | SOP12 | C x T | | 1043626 | 44709 | 0.043 |
| KCK1 | A25 | C x T | | 107586 | 4702 | 0.044 |
| KCK1 | ICE10 | C x T | | 935428 | 40901 | 0.044 |
| KCK1 | LVR | C x T | | 2074880 | 92339 | 0.045 |
| KCK1 | SAB | C x T | | 1159386 | 49588 | 0.043 |
| KCK1 | SAB19 | C x T | | 893729 | 39593 | 0.044 |
| KCK1 | SOP12 | C x T | | 1210030 | 53039 | 0.044 |
| PAG2 | A25 | C x T | | 78468 | 3439 | 0.044 |
| PAG2 | ICE10 | C x T | | 634316 | 28411 | 0.045 |
| PAG2 | LVR | C x T | | 1367191 | 61742 | 0.045 |

| | | | | | | |
|---|---|---|---|---|---|---|
| PAG2 | SAB | C x T | | 779241 | 33751 | 0.043 |
| PAG2 | SAB19 | C x T | | 602992 | 26812 | 0.044 |
| PAG2 | SOP12 | C x T | | 811659 | 36229 | 0.045 |
| TWN36 | A25 | C x T | | 97548 | 4230 | 0.043 |
| TWN36 | ICE10 | C x T | | 834223 | 36881 | 0.044 |
| TWN36 | LVR | C x T | | 1824278 | 81433 | 0.045 |
| TWN36 | SAB | C x T | | 1031616 | 44284 | 0.043 |
| TWN36 | SAB19 | C x T | | 790395 | 35220 | 0.045 |
| TWN36 | SOP12 | C x T | | 1076907 | 47770 | 0.044 |
| UTC1 | A25 | C x T | | 70457 | 2958 | 0.042 |
| UTC1 | ICE10 | C x T | | 571105 | 25413 | 0.044 |
| UTC1 | LVR | C x T | | 1226757 | 55219 | 0.045 |
| UTC1 | SAB | C x T | | 700213 | 30371 | 0.043 |
| UTC1 | SAB19 | C x T | | 543723 | 24187 | 0.044 |
| UTC1 | SOP12 | C x T | | 736565 | 32717 | 0.044 |
| UTC2 | A25 | C x T | | 74417 | 3197 | 0.043 |
| UTC2 | ICE10 | C x T | | 602891 | 26806 | 0.044 |
| UTC2 | LVR | C x T | | 1296909 | 58507 | 0.045 |
| UTC2 | SAB | C x T | | 743663 | 32260 | 0.043 |
| UTC2 | SAB19 | C x T | | 573713 | 25668 | 0.045 |
| UTC2 | SOP12 | C x T | | 776483 | 34557 | 0.045 |
| NOR511 | A25 | M x T | | 122624 | 5226 | 0.043 |
| NOR511 | ICE10 | M x T | | 1139922 | 49550 | 0.043 |
| NOR511 | LVR | M x T | | 2603770 | 113778 | 0.044 |
| NOR511 | SAB | M x T | | 1425235 | 60643 | 0.043 |
| NOR511 | SAB19 | M x T | | 1090892 | 47409 | 0.043 |
| NOR511 | SOP12 | M x T | | 1484611 | 64642 | 0.044 |
| NOR523 | A25 | M x T | | 112815 | 4883 | 0.043 |
| NOR523 | ICE10 | M x T | | 1032201 | 44986 | 0.044 |
| NOR523 | LVR | M x T | | 2322219 | 101896 | 0.044 |
| NOR523 | SAB | M x T | | 1285273 | 54622 | 0.042 |
| NOR523 | SAB19 | M x T | | 983031 | 42622 | 0.043 |
| NOR523 | SOP12 | M x T | | 1342275 | 58638 | 0.044 |
| UNP12 | A25 | M x T | | 101119 | 4254 | 0.042 |
| UNP12 | ICE10 | M x T | | 895121 | 37593 | 0.042 |
| UNP12 | LVR | M x T | | 1980730 | 83730 | 0.042 |
| UNP12 | SAB | M x T | | 1109875 | 45479 | 0.041 |
| UNP12 | SAB19 | M x T | | 851447 | 36084 | 0.042 |
| UNP12 | SOP12 | M x T | | 1160148 | 48941 | 0.042 |
| AHQT1 | CACG | G x G | | 3416204 | 148002 | 0.043 |
| AHQT1 | MED84 | G x G | | 3361660 | 187971 | 0.056 |
| AHQT1 | SLP | G x G | | 3328696 | 189792 | 0.057 |
| CACG | MED84 | G x G | | 3570377 | 193520 | 0.054 |
| CACG | SLP | G x G | | 3528904 | 197570 | 0.056 |
| MED84 | SLP | G x G | | 3500980 | 164704 | 0.047 |

| | | | | | | |
|---|---|---|---|---|---|---|
| CACN | NHN | N x N | | 3554430 | 48909 | 0.014 |
| AHQT1 | CACN | G x N | | 3349682 | 195561 | 0.058 |
| AHQT1 | NHN | G x N | | 3324482 | 193062 | 0.058 |
| CACG | CACN | G x N | | 3590384 | 170754 | 0.048 |
| CACG | NHN | G x N | | 3551846 | 172971 | 0.049 |
| MED84 | CACN | G x N | | 3523489 | 177903 | 0.05 |
| MED84 | NHN | G x N | | 3491227 | 176917 | 0.051 |
| SLP | CACN | G x N | | 3483131 | 185615 | 0.053 |
| SLP | NHN | G x N | | 3452238 | 183961 | 0.053 |
| AHQT1 | GAB1 | G x C | | 730827 | 45239 | 0.062 |
| AHQT1 | GAB2 | G x C | | 1682290 | 102780 | 0.061 |
| AHQT1 | KCK1 | G x C | | 1981058 | 123066 | 0.062 |
| AHQT1 | PAG2 | G x C | | 1312094 | 81888 | 0.062 |
| AHQT1 | TWN36 | G x C | | 1746066 | 109325 | 0.063 |
| AHQT1 | UTC1 | G x C | | 1171205 | 73499 | 0.063 |
| AHQT1 | UTC2 | G x C | | 1241732 | 77929 | 0.063 |
| CACG | GAB1 | G x C | | 760699 | 47408 | 0.062 |
| CACG | GAB2 | G x C | | 1764329 | 108089 | 0.061 |
| CACG | KCK1 | G x C | | 2082881 | 130312 | 0.063 |
| CACG | PAG2 | G x C | | 1373288 | 86595 | 0.063 |
| CACG | TWN36 | G x C | | 1830691 | 115224 | 0.063 |
| CACG | UTC1 | G x C | | 1231268 | 77515 | 0.063 |
| CACG | UTC2 | G x C | | 1301649 | 82176 | 0.063 |
| MED84 | GAB1 | G x C | | 750439 | 47567 | 0.063 |
| MED84 | GAB2 | G x C | | 1737986 | 109016 | 0.063 |
| MED84 | KCK1 | G x C | | 2052312 | 131200 | 0.064 |
| MED84 | PAG2 | G x C | | 1354279 | 86877 | 0.064 |
| MED84 | TWN36 | G x C | | 1803934 | 115526 | 0.064 |
| MED84 | UTC1 | G x C | | 1212922 | 78315 | 0.065 |
| MED84 | UTC2 | G x C | | 1284219 | 82746 | 0.064 |
| SLP | PAG2 | G x C | | 1345396 | 86463 | 0.064 |
| SLP | GAB1 | G x C | | 745670 | 47215 | 0.063 |
| SLP | GAB2 | G x C | | 1725197 | 108074 | 0.063 |
| SLP | KCK1 | G x C | | 2036038 | 130234 | 0.064 |
| SLP | TWN36 | G x C | | 1789942 | 114991 | 0.064 |
| SLP | UTC1 | G x C | | 1203888 | 77469 | 0.064 |
| SLP | UTC2 | G x C | | 1273984 | 82306 | 0.065 |
| AHQT1 | NOR511 | G x M | | 2480788 | 157418 | 0.063 |
| AHQT1 | NOR523 | G x M | | 2216473 | 141265 | 0.064 |
| AHQT1 | UNP12 | G x M | | 1894887 | 115160 | 0.061 |
| CACG | NOR511 | G x M | | 2615338 | 166314 | 0.064 |
| CACG | NOR523 | G x M | | 2331722 | 149064 | 0.064 |
| CACG | UNP12 | G x M | | 1986719 | 121191 | 0.061 |
| MED84 | NOR511 | G x M | | 2575152 | 166881 | 0.065 |
| MED84 | NOR523 | G x M | | 2296411 | 149358 | 0.065 |

| | | | | | | |
|---|---|---|---|---|---|---|
| MED84 | UNP12 | G x M | | 1959497 | 122176 | 0.062 |
| SLP | NOR511 | G x M | | 2558097 | 165848 | 0.065 |
| SLP | NOR523 | G x M | | 2281789 | 148129 | 0.065 |
| SLP | UNP12 | G x M | | 1947407 | 121339 | 0.062 |
| AHQT1 | A25 | G x T | | 148261 | 8524 | 0.057 |
| AHQT1 | ICE10 | G x T | | 1396059 | 84872 | 0.061 |
| AHQT1 | LVR | G x T | | 3334869 | 207273 | 0.062 |
| AHQT1 | SAB | G x T | | 1746019 | 106372 | 0.061 |
| AHQT1 | SAB19 | G x T | | 1343681 | 82582 | 0.061 |
| AHQT1 | SOP12 | G x T | | 1820733 | 110965 | 0.061 |
| CACG | A25 | G x T | | 152530 | 8879 | 0.058 |
| CACG | ICE10 | G x T | | 1462769 | 89166 | 0.061 |
| CACG | LVR | G x T | | 3571672 | 221848 | 0.062 |
| CACG | SAB | G x T | | 1830294 | 111735 | 0.061 |
| CACG | SAB19 | G x T | | 1413398 | 87674 | 0.062 |
| CACG | SOP12 | G x T | | 1911775 | 116626 | 0.061 |
| MED84 | A25 | G x T | | 150840 | 8851 | 0.059 |
| MED84 | ICE10 | G x T | | 1444032 | 89288 | 0.062 |
| MED84 | LVR | G x T | | 3498051 | 220313 | 0.063 |
| MED84 | SAB | G x T | | 1807001 | 111857 | 0.062 |
| MED84 | SAB19 | G x T | | 1393434 | 87345 | 0.063 |
| MED84 | SOP12 | G x T | | 1885911 | 116496 | 0.062 |
| SLP | A25 | G x T | | 149701 | 8783 | 0.059 |
| SLP | ICE10 | G x T | | 1432212 | 88588 | 0.062 |
| SLP | LVR | G x T | | 3461730 | 217689 | 0.063 |
| SLP | SAB | G x T | | 1792636 | 111337 | 0.062 |
| SLP | SAB19 | G x T | | 1380936 | 86550 | 0.063 |
| SLP | SOP12 | G x T | | 1871832 | 115994 | 0.062 |
| CACN | GAB1 | N x C | | 751709 | 47408 | 0.063 |
| CACN | GAB2 | N x C | | 1745259 | 109145 | 0.063 |
| CACN | KCK1 | N x C | | 2060090 | 130991 | 0.064 |
| CACN | PAG2 | N x C | | 1357932 | 86984 | 0.064 |
| CACN | TWN36 | N x C | | 1810024 | 116139 | 0.064 |
| CACN | UTC1 | N x C | | 1216126 | 78151 | 0.064 |
| CACN | UTC2 | N x C | | 1287524 | 82629 | 0.064 |
| NHN | GAB1 | N x C | | 748160 | 47344 | 0.063 |
| NHN | GAB2 | N x C | | 1733297 | 108370 | 0.063 |
| NHN | KCK1 | N x C | | 2044043 | 130055 | 0.064 |
| NHN | PAG2 | N x C | | 1348265 | 86428 | 0.064 |
| NHN | TWN36 | N x C | | 1796888 | 115252 | 0.064 |
| NHN | UTC1 | N x C | | 1207186 | 77584 | 0.064 |
| NHN | UTC2 | N x C | | 1278836 | 82111 | 0.064 |
| CACN | NOR511 | N x M | | 2589288 | 167347 | 0.065 |
| CACN | NOR523 | N x M | | 2309461 | 149560 | 0.065 |

| | | | | | | |
|---|---|---|---|---|---|---|
| CACN | UNP12 | N x M | | 1969078 | 122719 | 0.062 |
| NHN | NOR511 | N x M | | 2569555 | 165887 | 0.065 |
| NHN | NOR523 | N x M | | 2292474 | 148195 | 0.065 |
| NHN | UNP12 | N x M | | 1955006 | 121669 | 0.062 |
| CACN | A25 | N x T | | 3554430 | 48909 | 0.014 |
| CACN | ICE10 | N x T | | 1447148 | 89359 | 0.062 |
| CACN | LVR | N x T | | 3512330 | 220243 | 0.063 |
| CACN | SAB | N x T | | 1813000 | 111797 | 0.062 |
| CACN | SAB19 | N x T | | 1398633 | 87410 | 0.062 |
| CACN | SOP12 | N x T | | 1890361 | 116666 | 0.062 |
| NHN | A25 | N x T | | 149589 | 8831 | 0.059 |
| NHN | ICE10 | N x T | | 1435901 | 88500 | 0.062 |
| NHN | LVR | N x T | | 3477790 | 217948 | 0.063 |
| NHN | SAB | N x T | | 1800013 | 110997 | 0.062 |
| NHN | SAB19 | N x T | | 1388316 | 86634 | 0.062 |
| NHN | SOP12 | N x T | | 1876577 | 115751 | 0.062 |
| DENT | AHQT1 | D x G | | 1152795 | 86528 | 0.075 |
| DENT | CACG | D x G | | 1187840 | 89586 | 0.075 |
| DENT | MED84 | D x G | | 1180682 | 92151 | 0.078 |
| DENT | SLP | D x G | | 1176385 | 91810 | 0.078 |
| CACN | DENT | D x N | | 1179364 | 92160 | 0.078 |
| DENT | NHN | D x N | | 1171388 | 91453 | 0.078 |
| DENT | GAB1 | D x C | | 273822 | 18870 | 0.069 |
| DENT | GAB2 | D x C | | 637372 | 44037 | 0.069 |
| DENT | KCK1 | D x C | | 749612 | 52618 | 0.07 |
| DENT | PAG2 | D x C | | 512725 | 36198 | 0.071 |
| DENT | TWN36 | D x C | | 665863 | 47162 | 0.071 |
| DENT | UTC1 | D x C | | 458721 | 32477 | 0.071 |
| DENT | UTC2 | D x C | | 481807 | 34237 | 0.071 |
| DENT | NOR511 | D x M | | 930364 | 65313 | 0.07 |
| DENT | NOR523 | D x M | | 839748 | 59184 | 0.07 |
| DENT | UNP12 | D x M | | 723134 | 49397 | 0.068 |
| DENT | A25 | D x T | | 63920 | 4190 | 0.066 |
| DENT | ICE10 | D x T | | 531642 | 37223 | 0.07 |
| DENT | LVR | D x T | | 1179931 | 83547 | 0.071 |
| DENT | SAB | D x T | | 656016 | 45578 | 0.069 |
| DENT | SAB19 | D x T | | 508440 | 35781 | 0.07 |
| DENT | SOP12 | D x T | | 687169 | 48051 | 0.07 |

| | | | | | | | | | |
|---|---|---|---|---|---|---|---|---|---|
| **C x C**<br>$N_{cross} = 43$<br>$N_{total} = 116$ | 167.97 ±<br>10.73 | | | | | | | | |
| **C x M**<br>$N_{cross} = 10$<br>$N_{total} = 22$ | 9.16e-05<br>0.097<br>1.0 | 170.59 ±<br>25.03 | | | | | | | |
| **M x C**<br>$N_{cross} = 12$<br>$N_{total} = 25$ | 2.79e-03<br>4.81<br>0.0001 | -2.69e-03<br>-2.79<br>0.12 | 315.60 ±<br>43.44 | | | | | | |
| **M x M**<br>$N_{cross} = 4$<br>$N_{total} = 7$ | 2.88e-03<br>3.24<br>0.032 | 2.78e-03<br>2.37<br>0.30 | 9.03e-05<br>0.10<br>1.0 | 324.86 ±<br>84.50 | | | | | |
| **M x T**<br>$N_{cross} = 8$<br>$N_{total} = 14$ | 1.14e-03<br>1.19<br>0.96 | 1.05e-3<br>0.85<br>1.0 | -1.64e-03<br>-1.66<br>0.77 | -1.73e-03<br>-1.45<br>0.88 | 207.93 ±<br>38.29 | | | | |
| **T x M**<br>$N_{cross} = 7$<br>$N_{total} = 14$ | 4.33e-04<br>0.40<br>1.0 | 3.41e-04<br>0.26<br>1.0 | -2.35e-03<br>-2.13<br>0.45 | -2.44e-03<br>-1.89<br>0.62 | 7.11e-04<br>0.53<br>1.0 | 181.14 ±<br>3.32 | | | |
| **T x T**<br>$N_{cross} = 14$<br>$N_{total} = 37$ | 2.13e-04<br>0.28<br>1.0 | 1.21e-04<br>0.11<br>1.0 | -2.57e-03<br>-3.28<br>0.028 | -2.66e-03<br>-2.58<br>0.19 | -9.32e-04<br>-0.85<br>1.0 | -2.20e-04<br>-0.18<br>1.0 | 174.19 ±<br>19.71 | | |
| **T x C**<br>$N_{cross} = 25$<br>$N_{total} = 75$ | 1.49e-03<br>2.87<br>0.09 | -1.40e-03<br>-1.50<br>-0.85 | -1.29e-03<br>-2.30<br>0.34 | 1.38e-03<br>1.58<br>0.82 | -3.46e-04<br>-0.36<br>1.0 | -1.06e-03<br>-0.98<br>0.99 | -1.28e-03<br>-1.73<br>0.73 | 224.05 ±<br>17.80 | |
| **C x T**<br>$N_{cross} = 24$<br>$N_{total} = 60$ | -1.81e-04<br>-0.27<br>1.0 | -2.73e-04<br>-0.27<br>1.0 | -2.97e-03<br>-4.25<br>0.0007 | -3.06e-03<br>-3.15<br>0.043 | 1.33e-03<br>1.28<br>0.94 | -6.14e-04<br>-0.53<br>1.0 | 3.94e-04<br>0.4<br>1.0 | -1.27e-03<br>-2.57<br>0.20 | 163 ±<br>14.48 |
| | **C x C** | **C x M** | **M x C** | **M x M** | **M x T** | **T x M** | **T x T** | **T x C** | **C x T** |

| | Likelihood Ratio $\chi^2$ | *df* | *p* |
|---|---|---|---|
| Maternal Species | 21.80 | 2 | 1.84e-05 |
| Paternal Species | 0.10 | 2 | 0.95 |
| Maternal*Paternal | 2.81 | 4 | 0.59 |

| | | | |
|---|---|---|---|
| **C**<br>$N_{cross} = 77$<br>$N_{total} = 198$ | 167.13<br>± 10.12 | | |
| **M**<br>$N_{cross} = 24$<br>$N_{total} = 46$ | 2.30e-03<br>4.12<br><0.0001 | 271.34 ±<br>31.18 | |
| **T**<br>$N_{cross} = 46$<br>$N_{total} = 127$ | 7.42e-04<br>1.34<br>0.373 | 1.56e-03<br>-2.61<br>0.024 | 190.78<br>± 15.24 |
| | **C** | **M** | **T** |

Table S10. Pairwise differences in seed set per fruit assessed using a post-hoc Tukey method. Cross types involved *M. caespitosa* (C), *M. minor* (M), and *M. tilingii* (T), with the maternal parent in each cross listed first. $N_{cross}$ = number of unique maternal family combinations per cross type, and $N_{total}$ = total number of fruits scored per cross type. Values on diagonal are lsmeans +/- standard error. In each box below the diagonal, the uppermost value is the model estimate, the middle value is the z-ratio, and the bottom value is the *P*-value. Upper right: GLM type III ANOVA results of intra- and interspecific seed set with likelihood-ratio $\chi^2$ values for "Maternal Species" and "Paternal Species" (fixed effects) and "Maternal*Paternal" species interaction effect. Below ANOVA table: pairwise differences driven by maternal species. Shades of light gray denotes a $P < 0.05$, medium gray denotes a $P < 0.01$, and dark gray denotes a $P < 0.001$.

| | C x C | C x M | M x C | M x M | M x T | T x M | T x T | T x C | C x T |
|---|---|---|---|---|---|---|---|---|---|
| **C x C**<br>$N_{cross}$ = 43<br>$N_{total}$ = 115 | 0.95 ± 0.02 | | | | | | | | |
| **C x M**<br>$N_{cross}$ = 10<br>$N_{total}$ = 22 | 0.79<br>-0.68<br>1.0 | 0.96±0.02 | | | | | | | |
| **M x C**<br>$N_{cross}$ = 12<br>$N_{total}$ = 25 | 3.95<br>1.77<br>0.70 | 0.2<br>-1.90<br>0.62 | 0.84±0.09 | | | | | | |
| **M x M**<br>$N_{cross}$ = 4<br>$N_{total}$ =7 | 1.71<br>0.64<br>1.0 | 2.16<br>0.98<br>0.99 | 0.43<br>-2.46<br>0.25 | 0.92±0.05 | | | | | |
| **M x T**<br>$N_{cross}$ = 8<br>$N_{total}$ = 14 | 1.86e03<br>9.02<br><0.0001 | 2.35e03<br>8.89<br><0.0001 | 471.87<br>19.86<br><0.0001 | 1.09e03<br>17.41<br><0.0001 | 0.01±0.01 | | | | |
| **T x M**<br>$N_{cross}$ = 7<br>$N_{total}$ = 14 | 6.67<br>2.74<br>0.13 | 8.40<br>3.48<br>0.015 | 1.69<br>0.58<br>1.0 | 3.90<br>1.61<br>0.80 | 279.47<br>6.08<br><0.0001 | 0.76 ±0.10 | | | |
| **T x T**<br>$N_{cross}$ = 14<br>$N_{total}$ = 36 | 2.04<br>1.08<br>0.98 | 2.57<br>1.33<br>0.92 | 0.52<br>-0.75<br>1.0 | 1.19<br>0.19<br>1.0 | 1.1e-03<br>-7.97<br><0.0001 | 0.31<br>-3.27<br>0.03 | 0.91±0.04 | | |
| **T x C**<br>$N_{cross}$ = 25<br>$N_{total}$ = 75 | 2.15<br>1.31<br>0.94 | 0.37<br>-1.44<br>0.88 | 0.55<br>-0.72<br>1.0 | 0.79<br>-0.25<br>1.0 | 865.37<br>7.58<br><0.0001 | 3.10<br>3.37<br>0.022 | 0.95<br>-0.21<br>1.0 | 0.91±0.04 | |
| **C x T**<br>$N_{cross}$ = 24<br>$N_{total}$ = 60 | 85.65<br>16.99<br><0.0001 | 107.91<br>12.82<br><0.0001 | 21.68<br>3.76<br>0.005 | 50.03<br>4.57<br>0.0002 | 21.76<br>3.88<br>0.003 | 12.84<br>3.62<br>0.009 | 0.02<br>-6.15<br><0.0001 | 39.77<br>5.58<br><0.0001 | 0.20±0.07 |

| | Wald $\chi^2$ | *df* | *p* |
|---|---|---|---|
| Maternal Species | 3.76 | 2 | 0.15 |
| Paternal Species | 321.12 | 2 | < 2.2e-16 |
| Maternal*Paternal | 5848.23 | 4 | < 2.2e-16 |

Table S11. Pairwise differences in seed viability (morphological seed assessment per fruit) assessed using a post-hoc Tukey method. Cross types involved *M. caespitosa* (C), *M. minor* (M), and *M. tilingii* (T), with the maternal parent in each cross listed first. $N_{cross}$ = number of unique maternal family combinations per cross type, and $N_{total}$ = total number of fruits scored per cross type. Values on diagonal are lsmeans +/- standard error. In each box below the diagonal, the uppermost value is the model estimate, the middle value is the z-ratio, and the bottom value is the *P*-value. Upper right corner: GLMM type III ANOVA of intra- and interspecific morphological seed viability with Wald $\chi^2$ values for "Maternal Species" and "Paternal Species" (fixed effects) and "Maternal*Paternal" species interaction effect. Shades of light gray denotes a *P* <0.05, medium gray denotes a *P* <0.01, and dark gray denotes a *P* <0.001.

| | | | | | | | | | |
|---|---|---|---|---|---|---|---|---|---|
| **C x C**<br>$N_{cross}$ = 36<br>$N_{total}$ = 54 | 0.59 ± 0.06 | | | | | | | | |
| **C x M**<br>$N_{cross}$ = 8<br>$N_{total}$ = 13 | 3.06<br>3.53<br>0.012 | 0.32±0.07 | | | | | | | |
| **M x C**<br>$N_{cross}$ = 7<br>$N_{total}$ =13 | 1.70<br>1.46<br>0.88 | 1.80<br>1.22<br>0.95 | 0.46±0.09 | | | | | | |
| **M x M**<br>$N_{cross}$ = 3<br>$N_{total}$ =3 | 1.97<br>1.4<br>0.9 | 0.64<br>-1.20<br>0.96 | 1.16<br>0.45<br>1.0 | 0.42±0.1 | | | | | |
| **M x T**<br>$N_{cross}$ = 7<br>$N_{total}$ = 9 | 707.24<br>9.87<br><0.0001 | 230.8<br>7.71<br><0.0001 | 415.65<br>10.8<br><0.0001 | 359.88<br>9.7<br><0.0001 | 0±0 | | | | |
| **T x M**<br>$N_{cross}$ = 7<br>$N_{total}$ = 11 | 0.94<br>-0.15<br>1.0 | 0.031<br>-4.06<br>0.0016 | 0.55<br>-1.178<br>0.96 | 0.47<br>-1.85<br>0.65 | 752.73<br>9.18<br><0.0001 | 0.6±0.09 | | | |
| **T x T**<br>$N_{cross}$ = 10<br>$N_{total}$ = 19 | 0.53<br>-1.70<br>0.75 | 0.17<br>-3.96<br>0.0025 | 0.31<br>-2.53<br>0.22 | 0.27<br>-2.5<br>0.22 | 7.0e-04<br>-11.31<br><0.0001 | 0.56<br>-1.68<br>0.76 | 0.73±0.06 | | |
| **T x C**<br>$N_{cross}$ = 20<br>$N_T$ = 40 | 0.46<br>-2.75<br>0.13 | 6.72<br>4.48<br>0.0003 | 0.27<br>-3.35<br>0.023 | 4.31<br>2.89<br>0.09 | 1.5e04<br>10.78<br><0.0001 | 2.06<br>2.28<br>0.36 | 1.16<br>0.58<br>1.0 | 0.76±0.05 | |
| **C x T**<br>$N_{cross}$ = 23<br>$N_{total}$ = 47 | 4.21<br>5.82<br><0.0001 | 1.37<br>0.93<br>0.99 | 2.47<br>2.06<br>0.5 | 2.14<br>1.52<br>0.85 | 168.06<br>8.28<br><0.0001 | 4.48<br>3.38<br>0.021 | 0.13<br>-7.23<br><0.0001 | 9.23<br>5.9<br><0.0001 | 0.25 ±0.05 |
| | **C x C** | **C x M** | **M x C** | **M x M** | **M x T** | **T x M** | **T x T** | **T x C** | **C x T** |

| | Wald $\chi^2$ | *df* | *p* |
|---|---|---|---|
| Maternal Species | 13.07 | 2 | 1.46e-3 |
| Paternal Species | 37.75 | 2 | 6.36e-09 |
| Maternal*Paternal | 680.30 | 4 | < 2.2e-16 |

Table S12. Pairwise differences in seed viability (germination rate per fruit) assessed using a post-hoc Tukey method. Cross types involved *M. caespitosa* (C), *M. minor* (M), and *M. tilingii* (T), with the maternal parent in each cross listed first. $N_{cross}$ = number of unique maternal family combinations per cross type, and $N_{total}$ = total number of fruits scored per cross type. Values on diagonal are lsmeans +/- standard error. In each box below the diagonal, the uppermost value is the model estimate, the middle value is the z-ratio, and the bottom value is the *P*-value. Upper right corner: GLMM type III ANOVA of intra- and interspecific germination rate with Wald $\chi^2$ values for "Maternal Species" and "Paternal Species" (fixed effects) and "Maternal*Paternal" species interaction effect. Shades of light gray denotes a *P* <0.05, medium gray denotes a *P* <0.01, and dark gray denotes a *P* <0.001.

| | | | | | | | | | |
|---|---|---|---|---|---|---|---|---|---|
| **C x C**<br>$N_{cross}$=17<br>$N_{total}$=199 | 49.15 ±<br>2.04 | | | | | | | | |
| **C x M**<br>$N_{cross}$= 5<br>$N_{total}$= 57 | 1.36<br>4.51<br>0.0002 | 36.11 ±<br>2.41 | | | | | | | |
| **M x C**<br>$N_{cross}$= 4<br>$N_{total}$= 56 | 1.14<br>1.97<br>0.56 | 1.20<br>1.96<br>0.57 | 43.25 ±<br>2.76 | | | | | | |
| **M x M**<br>$N_{cross}$ = 2<br>$N_{total}$= 23 | 1.16<br>1.60<br>0.81 | 0.85<br>-2.22<br>0.39 | 1.02<br>0.29<br>1.0 | 42.31± 3.56 | | | | | |
| **M x T** | | | | | | | | | |
| **T x M**<br>$N_{cross}$ = 4<br>$N_{total}$= 54 | 1.11<br>1.29<br>0.94 | 0.82<br>-3.57<br>0.011 | 0.98<br>-0.22<br>1.0 | 0.96<br>-0.59<br>1.0 | | 44.19 ±<br>3.16 | | | |
| **T x T**<br>$N_{cross}$ = 8<br>$N_{total}$= 93 | 1.01<br>0.17<br>1.0 | 0.74<br>-3.35<br>0.023 | 0.89<br>-1.35<br>1.0 | 0.87<br>-1.35<br>0.92 | | 0.91<br>-1.23<br>0.95 | 48.56± 2.79 | | |
| **T x C**<br>$N_{cross}$= 43<br>$N_{total}$= 93 | 1.18<br>3.2<br>0.037 | 1.16<br>1.76<br>0.71 | 1.03<br>0.48<br>1.0 | 0.99<br>-0.12<br>1.0 | | 0.95<br>-0.79<br>1.0 | 0.86<br>-2.66<br>0.16 | 41.82 ± 2.09 | |
| **C x T**<br>$N_{cross}$ = 4<br>$N_{total}$ = 44 | 0.96<br>-0.76<br>1.0 | 0.70<br>-4.55<br>0.0002 | 084<br>-2.02<br>0.53 | 0.82<br>-1.93<br>0.59 | | 0.86<br>-1.67<br>0.77 | 1.06<br>0.92<br>0.99 | 0.82<br>-2.79<br>0.12 | 51.29 ±<br>2.76 |
| | **C x C** | **C x M** | **M x C** | **M x M** | **M x T** | **T x M** | **T x T** | **T x C** | **C x T** |

| | Wald $\chi^2$ | *df* | *p* |
|---|---|---|---|
| Maternal Species | 11.63 | 2 | 2.99e-03 |
| Paternal Species | 24.02 | 2 | 6.1e-06 |
| Maternal*Paternal | 93.40 | 3 | < 2.2e-16 |

Table S13. Pairwise differences in days to flower (from seedling) assessed using a post-hoc Tukey method. Cross types involved *M. caespitosa* (C), *M. minor* (M), and *M. tilingii* (T), with the maternal parent in each cross listed first. $N_{cross}$ = number of unique maternal family combinations per cross type, and $N_{total}$ = total number of individuals scored to flowering per cross type. Values on diagonal are lsmeans +/- standard error. In each box below the diagonal, the uppermost value is the model estimate, the middle value is the z-ratio, and the bottom value is the *P*-value. Upper right corner: GLMM type III ANOVA of intra- and interspecific days to first flower with Wald $\chi^2$ values for "Maternal Species" and "Paternal Species" (fixed effects) and "Maternal*Paternal" species interaction effect. Shades of light gray denotes a *P* <0.05, medium gray denotes a *P* <0.01, and dark gray denotes a *P* <0.001.

| | | | | | | | | | |
|---|---|---|---|---|---|---|---|---|---|
| **C x C**<br>$N_{cross}$ = 17<br>$N_{total}$ = 136 | 0.90 ± 0.02 | | | | | | | | |
| **C x M**<br>$N_{cross}$ = 5<br>$N_{total}$ = 38 | 177.91<br>16.0<br><0.0001 | 0.05 ± 0.01 | | | | | | | |
| **M x C**<br>$N_{cross}$ = 4<br>$N_{total}$ = 34 | 176.49<br>17.85<br><0.0001 | 1.01<br>0.02<br>1.0 | 0.05 ± 0.01 | | | | | | |
| **M x M**<br>$N_{cross}$ = 2<br>$N_{total}$ = 13 | 0.55<br>-1.39<br>0.90 | 3.1e-03<br>-20.16<br><0.0001 | 3.1e-03<br>-16.92<br><0.0001 | 0.94± 0.02 | | | | | |
| **M x T** | | | | | | | | | |
| **T x M**<br>$N_{cross}$ = 4<br>$N_{total}$ = 26 | 757.55<br>15.68<br><0.0001 | 4.26<br>5.07<br><0.0001 | 4.29<br>3.06<br>0.057 | 1.38e03<br>20.24<br><0.0001 | | 0.01 ± 0 | | | |
| **T x T**<br>$N_{cross}$ = 8<br>$N_{total}$ = 57 | 2.35<br>2.45<br>0.25 | 0.01<br>-10.16<br><0.0001 | 0.01<br>-10.47<br><0.0001 | 4.29<br>3.04<br>0.06 | | 3.1e-03<br>-14.35<br><0.0001 | 0.79 ± 0.05 | | |
| **T x C**<br>$N_{cross}$ = 9<br>$N_{total}$ = 65 | 14.69<br>12.26<br><0.0001 | 12.11<br>6.40<br><0.0001 | 0.08<br>-7.98<br><0.0001 | 0.04<br>-7.35<br><0.0001 | | 51.55<br>10.86<br><0.0001 | 0.16<br>-6.68<br><0.0001 | 0.38± 0.05 | |
| **C x T**<br>$N_{cross}$ = 5<br>$N_{total}$ = 29 | 15.77<br>9.98<br><0.0001 | 0.09<br>-6.57<br><0.0001 | 0.09<br>-6.06<br><0.0001 | 28.80<br>7.20<br><0.001 | | 0.02<br>-8.46<br><0.0001 | 0.15<br>-8.53<br><0.0001 | 1.07<br>0.20<br>1.0 | 0.36 ± 0.06 |
| | **C x C** | **C x M** | **M x C** | **M x M** | **M x T** | **T x M** | **T x T** | **T x C** | **C x T** |

| | Wald $\chi^2$ | *df* | *p* |
|---|---|---|---|
| Maternal Species | 376.71 | 2 | < 2.2e-16 |
| Paternal Species | 293.33 | 2 | < 2.2e-16 |
| Maternal*Paternal | 10162.86 | 3 | < 2.2e-16 |

Table S14. Pairwise differences in F1 pollen viability assessed using a post-hoc Tukey method. Cross types involved *M. caespitosa* (C), *M. minor* (M), and *M. tilingii* (T), with the maternal parent in each cross listed first. $N_{cross}$ = number of unique maternal family combinations per cross type$N_{total}$ = total number of flowers scored for viable pollen per cross type. Values on diagonal are lsmeans +/- standard error. In each box below the diagonal, the uppermost value is the model estimate, the middle value is the z-ratio, and the bottom value is the *P*-value. Upper right corner: GLMM type III ANOVA of F1 pollen viability with Wald $\chi^2$ values for "Maternal Species" and "Paternal Species" (fixed effects) and "Maternal*Paternal" species interaction effect. Shades of light gray denotes a *P* <0.05, medium gray denotes a *P* <0.01, and dark gray denotes a *P* <0.001.

| | | | | | | | | | |
|---|---|---|---|---|---|---|---|---|---|
| **C x C**<br>$N_{cross}$ = 17<br>$N_{total}$ = 76 | 206.03 ± 45.63 | | | | | | | | |
| **C x M**<br>$N_{cross}$ = 5<br>$N_{total}$ = 22 | 39<br>8.5<br><0.0001 | 5.28 ± 2.09 | | | | | | | |
| **M x C**<br>$N_{cross}$ = 4<br>$N_{total}$ = 17 | 65.01<br>16.42<br><0.0001 | 0.60<br>-1.02<br>0.98 | 3.17 ± 0.97 | | | | | | |
| **M x M**<br>$N_{cross}$ = 2<br>$N_{total}$ = 9 | 0.41<br>-1.89<br>0.62 | 0.01<br>-18.77<br><0.0001 | 6.25e-03<br>-11.36<br><0.0001 | 507.06± 214.34 | | | | | |
| **M x T** | | | | | | | | | |
| **T x M**<br>$N_{cross}$ = 4<br>$N_{total}$ = 11 | 42.3<br>7.97<br><0.0001 | 1.08<br>0.34<br>1.0 | 0.65<br>-0.84<br>1.0 | 104.11<br>16.58<br><0.0001 | | 4.87 ± 2.02 | | | |
| **T x T**<br>$N_{cross}$ = 8<br>$N_{total}$ = 34 | 1.23<br>0.57<br>1.0 | 0.03<br>-6.97<br><0.0001 | 0.02<br>-9.32<br><0.0001 | 3.04<br>2.15<br>0.44 | | 0.03<br>-7.46<br><0.0001 | 167± 49.58 | | |
| **T x C**<br>$N_{cross}$ = 9<br>$N_{total}$ = 43 | 4.47<br>8.80<br><0.0001 | 8.72<br>4.68<br>0.0001 | 0.07<br>-9.92<br><0.0001 | 0.09<br>-4.94<br><0.0001 | | 9.46<br>5.12<br><0.0001 | 0.28<br>-3.89<br>0.0032 | 46.09 ± 11.02 | |
| **C x T**<br>$N_{cross}$ = 4<br>$N_{total}$ = 19 | 8.34<br>6.39<br><0.0001 | 0.21<br>-3.30<br>0.027 | 0.13<br>-4.91<br><0.0001 | 20.53<br>5.92<br><0.0001 | | 0.2<br>-3.22<br>0.034 | 0.15<br>-10.66<br><0.0001 | 1.87<br>1.68<br>0.76 | 24.70 ± 7.06 |
| | **C x C** | **C x M** | **M x C** | **M x M** | **M x T** | **T x M** | **T x T** | **T x C** | **C x T** |

| | Wald $\chi^2$ | *df* | *p* |
|---|---|---|---|
| Maternal Species | 294.77 | 2 | < 2.2e-16 |
| Paternal Species | 90.35 | 2 | < 2.2e-16 |
| Maternal*Paternal | 4896.76 | 3 | < 2.2e-16 |

Table S15. Pairwise differences in F1 seed production assessed using a post-hoc Tukey method. Cross types involved *M. caespitosa* (C), *M. minor* (M), and *M. tilingii* (T), with the maternal parent in each cross listed first. $N_{cross}$ = number of unique maternal family combinations per cross type and $N_{total}$ = total number of fruits scored per cross type. Values on diagonal are lsmeans +/- standard error. In each box below the diagonal, the uppermost value is the model estimate, the middle value is the z-ratio, and the bottom value is the *P*-value. Upper right corner: GLMM type III ANOVA of F1 seed set Wald $\chi^2$ values for "Maternal Species" and "Paternal Species" (fixed effects) and "Maternal*Paternal" species interaction effect. Shades of light gray denotes a *P* <0.05, medium gray denotes a *P* <0.01, and dark gray denotes a *P* <0.001.
